# TriPort tangential flow filtration: a lab-assembled system for EV isolation across diverse cell-culture sources

**DOI:** 10.64898/2026.10.03.756465

**Authors:** Drishya Mainali, Mangesh D Hade, Anjie Ge, Hiba Ezzarouli, Simon Maksour, Lachlan Thompson, Cesar L Moreno, G Gregory Neely, Susannah Hallal, Claire Goldsbury, Setty Magaña, Laura Piccio

## Abstract

Tangential flow filtration (TFF) is an effective method for scalable EV isolation. Here, we describe and validate a reference protocol for TriPort-TFF, a laboratory-assembled TFF system featuring a three-port reservoir design enabling sample circulation, retentate return, and diafiltration. Constructed from individually sourced, commercially available components with detailed assembly and operating parameters, TriPort-TFF can be built at a lower cost than commercially integrated TFF platforms. We evaluated the system using conditioned media (CM) from human MDM cultured with EV-depleted human serum, human cerebral organoids in serum-free medium, and mouse BMDM cultured with EV-depleted foetal bovine serum (FBS) unstimulated or LPS-stimulated. The system reproducibly isolated EVs from all three sources, and biological replicates of BMDM showed distinct proteomic clusters between LPS-stimulated and unstimulated conditions. The requirement for subsequent size-exclusion chromatography (SEC) after TFF depended on the sample’s serum-presence. In serum-containing CM, TFF alone co-concentrated serum proteins alongside EVs, subsequent SEC improved the particle-to-protein ratio, reduced inter-replicate variability and improved proteomic depth. TFF alone in serum-free organoid CM already yielded higher particle-to-protein ratios than TFF-only preparations from serum-containing CM, with SEC further improving this ratio. TriPort-TFF offers an accessible system for EV-isolation that can be combined with SEC for diverse sample types.

## Introduction

Extracellular vesicles (EVs) are lipid bilayer-enclosed, cell-derived nanoparticles that harbor and transfer diverse bioactive cargo, including proteins, lipids, and nucleic acids, to recipient cells. Once regarded merely as cellular ‘debris’, EVs are now recognised as novel therapeutic vehicles and non-invasive biomarker reservoirs. As understanding of the complexity of EV biogenesis and biophysical heterogeneity has continued to evolve, so has the need for accessible, reproducible and validated EV isolation and characterization protocols (1).

The choice of EV isolation method crucially determines the purity and the biological interpretation of EV molecular signals. This is especially important for EVs that are isolated from serum-supplemented cell-culture systems, where abundant serum proteins typically co-isolate with EVs and can confound downstream analyses. Despite being a recognised problem in the field, there is a lack of standardised isolation approaches that effectively remove serum contaminants from EVs (2,3). Ultracentrifugation, size-exclusion chromatography (SEC), and polymer-based precipitation remain the most widely used EV isolation methods. Each offers a unique balance and trade-off between EV purity, yield, and scalability (4,5), underscoring current Minimal Information for Studies of Extracellular Vesicles (MISEV) recommendations that emphasise and encourage transparent reporting of EV isolation workflows rather than endorsing one universal method (1). The choice of an EV isolation method is ultimately informed by downstream applications, single or bulk-EV analysis and accessibility to appropriate equipment.

Tangential flow filtration (TFF) has emerged as a promising method for scalable EV isolation, shown to outperform ultracentrifugation in yield, reproducibility, processing time, and scalability (6,2). Compared to ultracentrifugation, TFF better maintains EV morphology and integrity, and can substantially improve EV purity when paired with SEC (2,7). In a TFF system, the sample flows parallel, or tangential to a porous membrane rather than directly into it. A uniform flow rate is maintained across the membrane, and particles smaller than its molecular weight cut-off pass through into the permeate, while larger particles are retained in the recirculating retentate (8). Due to the sample flow running parallel to the membrane surface, TFF continuously sweeps away any material that would otherwise accumulate and foul a static filter, allowing much larger sample volumes to be processed than a dead-end filtration. Repeated recirculation concentrates the retained particles, while diafiltration, the continuous addition of fresh buffer as permeate is removed, exchanges the surrounding medium and washes out smaller soluble impurities (2,6). TFF has been used to concentrate EVs from large volumes of biological fluids, and its operating principle, including flow rate and membrane fouling, have been characterised in detail in industrial and pharmaceutical bioprocessing contexts (8–10). Continuous TFF systems have also been applied beyond the concentration of cell culture, for example to non-mammalian sources such as dairy by-products, bacteria and yeast (3,11,12).

Despite these advantages, TFF remains uncommon in standard cell biology laboratories. This is not because scalable TFF options do not exist. Several companies offer systems that can be configured for a wide range of starting sample (2,7,13). However, these TFF systems are commercially integrated platforms that typically require laboratories to purchase all specific manufacturer’s hardware, cartridges and accessories to ensure system compatibility which creates a financial barrier to the adoption of TFF systems in standard laboratories. What is largely missing is a validated, fully specified protocol for a system that is constructed from individually-sourced, commercially-available components rather than a purchased platform, that can be readily assembled, modified, and maintained in any standard laboratory at a fraction of the cost (see Supplementary Table 6). Published TFF protocols for EV isolation illustrate this gap. Some studies report detailed operating parameters, flow rates and pressure readings for building commercial TFF systems that are typically inaccessible for standard laboratories due to sample volume, cost and scale (8–10). Other studies report laboratory-built TFF systems but lack the details needed to reproduce the system independently (7).

Here, we describe and validate TriPort-TFF, a laboratory-assembled three-port reservoir design system. Constructed from individually sourced, commercially available components, we describe the assembly and operating parameters in full. TriPort-TFF was tested across three biologically and compositionally distinct conditioned media (CM) sources from human and mouse macrophage or human cerebral organoid cell-cultures. We found that TriPort-TFF performed consistently on all sources of CM tested. Furthermore, we demonstrate that subsequent application of size exclusion chromatography to TFF-derived EV isolates is necessary to enrich the EV preparations by removing serum protein contaminants coming from serum-supplemented media preparations that become co-concentrated with EVs during the TFF protocol. Additionally, we provide a TFF membrane cleaning and reuse protocol that supports sustained, reproducible operation across repeated runs. In summary, our aim was to validate an accessible TFF protocol, applicable to diverse samples and that can be readily implemented and adapted to any laboratory, without reliance on industrial-grade TFF.

## Methods

### 1. Isolation, culture and EV harvest of human monocytes derived macrophages (MDM)

Blood samples (30ml) were collected by venipuncture from a healthy individual (n = 1) under informed consent. The blood samples were processed to separate human serum, and peripheral blood mononuclear cells (PBMCs) were isolated and cultured immediately after the draw as described below. All procedures were reviewed and approved by the University of Sydney Human Ethics Committee (Project No. 2026/HE000285).

For PBMC isolation, whole blood (30 ml) collected from a healthy individual was first diluted 1:1 with sterile Dulbecco’s Phosphate-Buffered Saline (dPBS; Gibco, 20012027). The blood was then carefully layered over 15 ml Ficoll-Paque (Cytiva, 17144003) in a 50 ml conical tube and centrifuged (1200 x g, 30 min, 22 °C; acceleration = 5, deceleration = 0). The peripheral blood mononuclear cells (PBMC) buffy coat at the Ficoll-plasma interphase were carefully removed with a sterile pipette and transferred to a fresh 50ml tube. The PBMCs were washed twice with sterile ice-cold PBS (500 x g, 10 min, 4°C; acceleration = 9, deceleration = 9). The PBMC pellet was resuspended with 10ml PBS, and the cell count and viability were determined with a hemocytometer.

The PBMCs were recentrifuged (400 x g, for 10 min, 4 °C) and seeded at a density of 5 million cells per well (30 million cells in 6-well plates) in RPMI medium supplemented with 1X sodium-pyruvate (Gibco, 11360070), 1X L-Glutamine (Gibco, 25030081), 1X penicillin-streptomycin (Gibco, 15140122), and 1X Non-Essential Amino Acids (Gibco, 11140050). The cells were incubated for 2 h at 37 °C to allow monocyte adherence on the culture plate. Non-adherent cells were removed, and the adhered cells were washed twice with PBS. The cells were cultured for 7 days with RPMI medium, supplemented with 5% heat-inactivated EV-depleted autologous human serum (EVdep-HuS) and 50ng/ml of macrophage colony-stimulating factor (m-CSF) (BioLegend, 574806), with the media refreshed at day 3.

30ml of conditioned media (CM) was collected at day 3 and 7 of the culture. The CM was clarified from cellular debris (1000 x g, 10 min, 4 °C) before freezing at -80 °C. To generate technical replicates of human MDM (n=3), day 3 and 7 media were pooled (total 60ml) and then split into three equal volumes of 20ml for EV processing (Figure 2).

#### Heat-inactivated EV-depleted Serum (EVdep-HuS)

Autologous serum was produced by centrifuging whole blood (1000 x g, 10 min, 22°C). The serum was then heat inactivated (55 °C, 30 mins). The heat inactivated serum was aliquoted into 1.5 ml sterile ultracentrifugation Eppendorf tubes and was subjected to 100, 000 x g for 2 h at 4 °C (Hitachi CP100NX; rotor P50A3) to obtain EV-depleted human serum (EVdep-HuS).

### 2. Mouse bone marrow-derived macrophages (BMDM) isolation, culture and stimulation

Mouse BMDM (n=3) were collected from 10-week-old female from C57BL/6 mice were cultured following previously described methods (14). The mouse sample used in this study is exempted from ethics approval under the University of Sydney Australian Ethics Committee (AEC) guidelines.

Briefly, mouse femur was collected immediately following euthanasia, with major muscles and connective tissues being removed. Epiphyses of femur was removed, and bone marrow was collected by centrifugation (6000 rpm, 3.5 min, room temperature (RT)) of each bone in a perforated 0.6 mL microcentrifuge tube nested within a 1.5 mL microcentrifuge tube, containing 50 µL DMEM in both inner and outer tubes. The supernatant was discarded and the marrow pellet was resuspended in 1 mL 1X red blood cell (RBC) lysis buffer for 3 min RT. This reaction is stopped by PBS top-up and centrifugation to collect cell pellets (550 xg, 5mins, 4°C). The bone marrow suspension was further washed in PBS in a 15 mL tube (550 × g, 5 min, 4 °C), resuspended in 2 mL DMEM, passed through a pre-wet 40µM cell strainer (15-1040, Biologix) into a fresh tube, then centrifuged (550 × g, 5 min, 4°C). Cells were resuspended in fresh DMEM and counted using trypan blue.

Bone marrow cells were seeded at 8.5 x10^5^/ml concentration (6 x 10^6^ cells altogether) in DMEM supplemented with 10% EV-depleted FBS and 45 ng/mL recombinant M-CSF to induce their differentiation into BMDMs. On the day 4 of culture, the media was refreshed, and then on the day 7, BMDMs were stimulated with 1 µg/mL Lipopolysaccharide (LPS) for 24 h. Following LPS stimulation, the CM was harvested (7 ml) from each plate and retained for TFF-based EV isolation. Adherent BMDMs were washed twice with PBS, followed by incubation with 5 ml TrypLE (warmed at 37 °C) for 10-15 min to detach cells. Detached cells were further dislodged by scraping, and washed twice with PBS (550 × g, 5 min, 4°C). The resulting cell pellet was resuspended in 1 mL PBS and counted using trypan blue.

CM from n=3 independent unstimulated (BMDM) and LPS-stimulated (BMDM_LPS) cultures (biological replicates from three independent mice, 7 ml each) were used for TFF-based EV isolation followed by SEC and post-SEC proteomic characterization.

#### Preparation of EV-depleted Fetal Bovine Serum (FBS)

Heat-inactivated FBS aliquots (25 mL each) were ultracentrifuged (100,000 × g for 16 h at 4°C; Hitachi CP100NX ultracentrifuge, P50AT2 rotor) to deplete endogenous bovine EVs. Following ultracentrifugation, the EV-depleted supernatant was carefully decanted, and the bottom ∼1 mL was left undisturbed to avoid disrupting the pelleted material. EV-depleted FBS aliquots were stored at -80 °C, with a working aliquot maintained at 4°C for immediate use in complete media preparation.

### 3. Human Cerebral Organoid

#### Stem cell maintenance

All procedures involving human induced pluripotent stem cell lines (iPSC) used in these studies were approved by the University of Sydney Human Ethics Committee (HREC 2020/417). The human iPSC line Scti003a (Stemcell Technologies 200-0511) was used to generate cerebral brain organoids. Cells were maintained on Matrigel (Merck CLS354277) coated plates in mTeSR1 or mTeSR-plus (Stemcell Technologies, 85857 or 100-1130, respectively). Cells were routinely passaged using RelesR (Stemcell technologies 100-0483). All cell lines were karyotyped before all experiments and were used for no more than 20 passages.

#### Organoid generation and media collection

Organoids were generated following standard protocols (Giandomenico, Sutcliffe, and Lancaster, 2021) using the STEMdiff Cerebral organoid kit (Stemcell technologies, 08570) (15). Briefly, 50-70% confluent stem cells were brought to a single cell suspension with 9000 cells plated per well into ultra-low attachment u-bottom plates with embryoid formation medium and 10 μM ROCK inhibitor (Day 0). The medium was refreshed every second day. On day 5, embryoid bodies were switched to induction medium. On day 7, organoids were embedded into Matrigel and cultured in expansion medium until day 10 when the medium was switched to Maturation medium for the remainder of the differentiation process. The organoids were cultured on a shaker on day 12 using low attachment plates, and media was refreshed 3 times per week throughout the maturation period. Media containing EVs were harvested from 12 cerebral organoids between day 90-100 (10ml of CM was collected at day 90, 93, 96, and 100). The harvested medium was clarified (1000 x g, 10 min, 4 °C) to remove cellular debris, and supernatant was stored at -80 °C before EV isolation.

### 4. Sample pre-processing

All harvested CM were thawed once at 4 °C overnight and processed immediately. The CM was centrifuged twice at 4000 x g, 20 min, 4 °C to remove larger cell debris, followed by filtration at 0.45 μm filter under sterile conditions (Figure 2). The sample was then diluted to final volumes of 35 ml with filtered sterile PBS for TFF.

### 5. Tangential Flow Filtration (TFF)

TriPort-TFF, a laboratory-assembled TFF system, was constructed from individually sourced, commercially available components using a peristaltic pump-driven hollow-fiber filtration design (Table 1, Figure 1.1A). The system is built around a custom three-port cap assembly (blunt needles and 3-way stopcocks) enabling sample circulation, retentate return, and diafiltration/buffer exchange. This system was validated using CM from three biologically distinct sources: (1) human monocyte-derived macrophages (Human MDM; n=3 technical replicates from one pooled conditioned-media batch), (2) human cerebral organoids (Organoid; n=2 technical replicates), and (3) mouse bone marrow-derived macrophages (BMDM; n=3 biological replicates from independent animals), assessed under unstimulated and LPS-stimulated conditions.

**Table 1:**
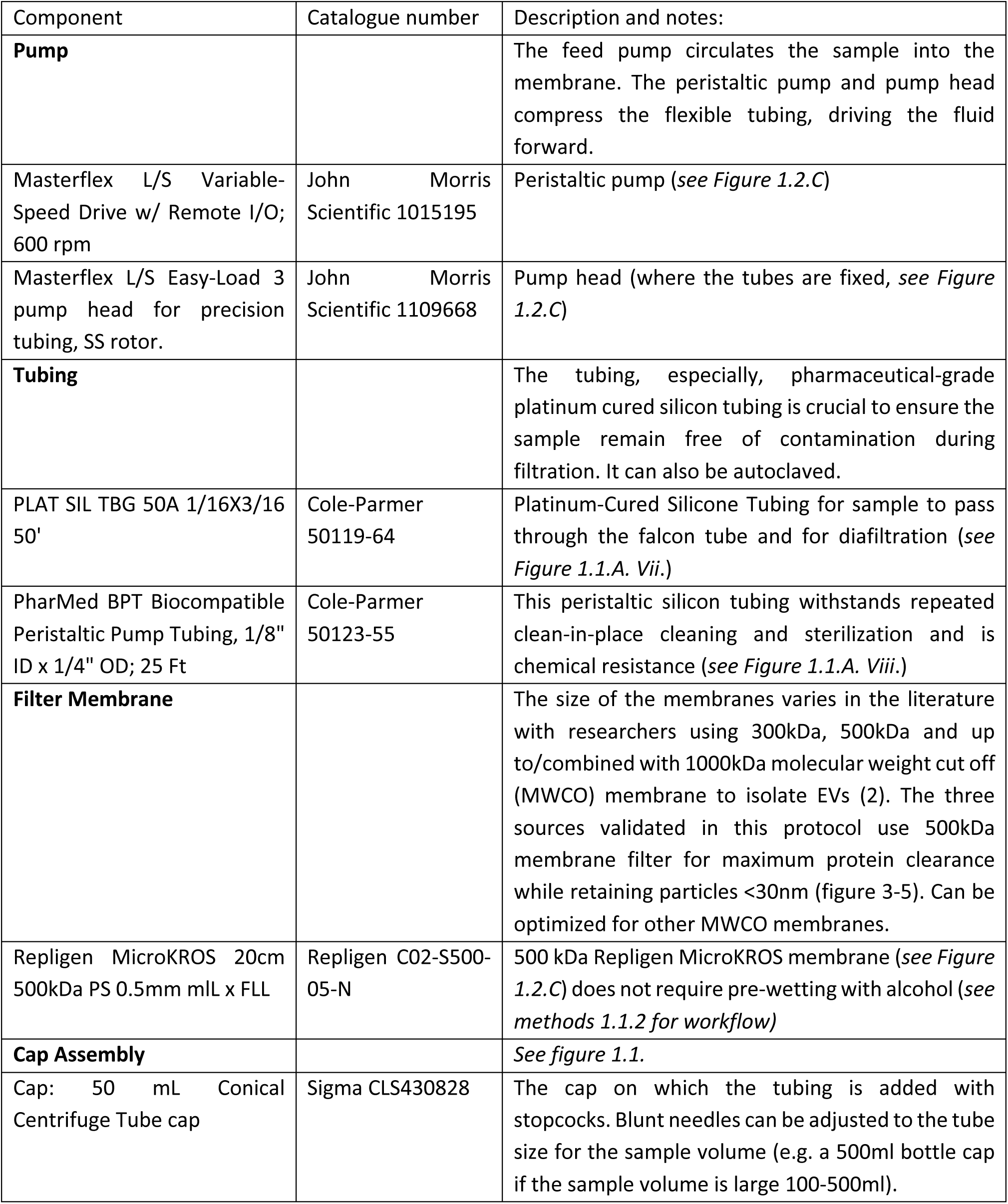
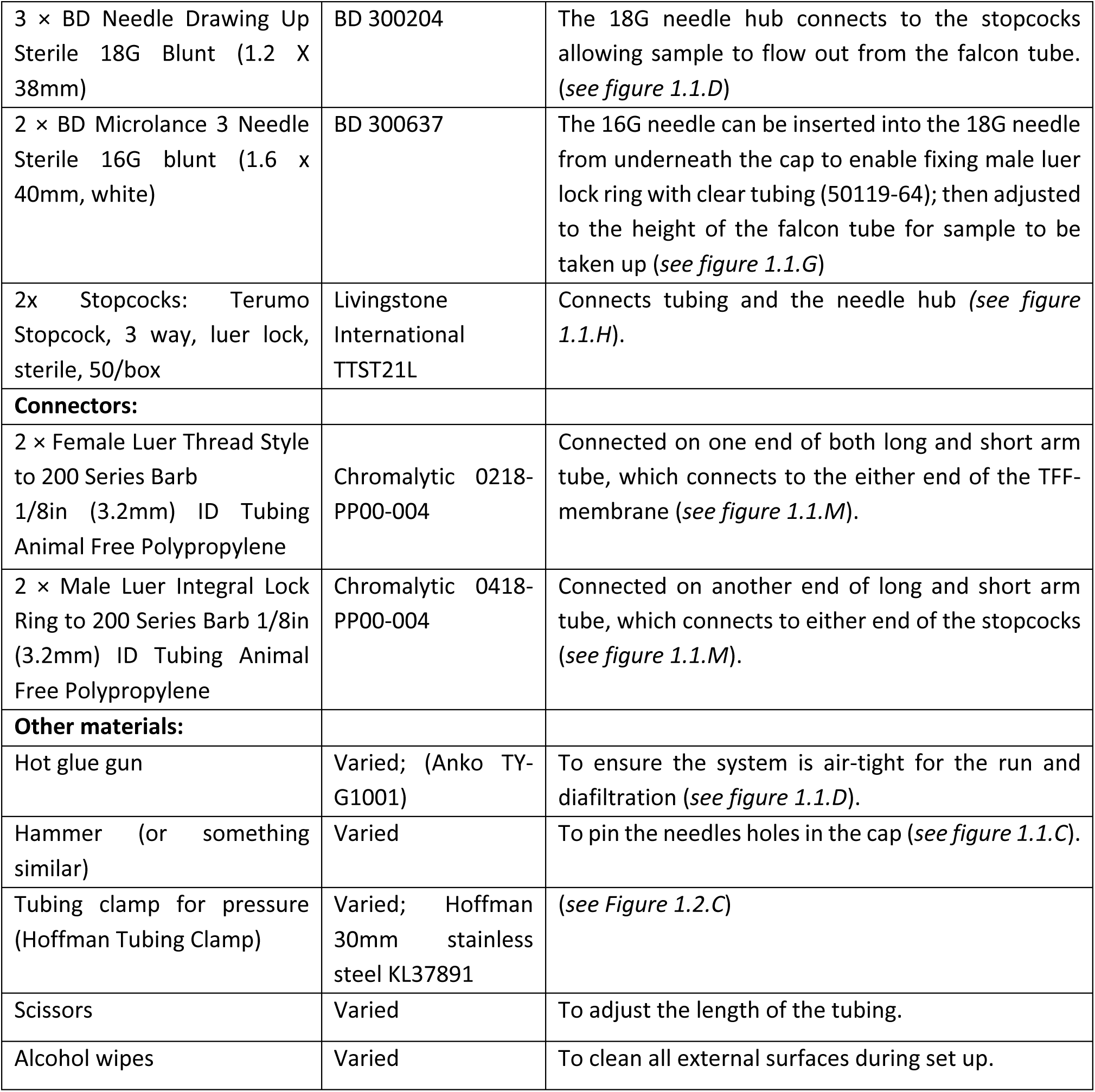
Materials, catalogue numbers, and notes for individually sourced components of the lab-assembled TFF system.

| Component | Catalogue number | Description and notes: |
| --- | --- | --- |
| **Pump** |  | The feed pump circulates the sample into the membrane. The peristaltic pump and pump head compress the flexible tubing, driving the fluid forward. |
| Masterflex L/S Variable-Speed Drive w/ Remote I/O; 600 rpm | John Morris Scientific 1015195 | Peristaltic pump (*see Figure 1.2.C*) |
| Masterflex L/S Easy-Load 3 pump head for precision tubing, SS rotor. | John Morris Scientific 1109668 | Pump head (where the tubes are fixed, *see Figure 1.2.C*) |
| **Tubing** |  | The tubing, especially, pharmaceutical-grade platinum cured silicon tubing is crucial to ensure the sample remain free of contamination during filtration. It can also be autoclaved. |
| PLAT SIL TBG 50A 1/16X3/16 50' | Cole-Parmer 50119-64 | Platinum-Cured Silicone Tubing for sample to pass through the falcon tube and for diafiltration (*see Figure 1.1.A. Vii.*) |
| PharMed BPT Biocompatible Peristaltic Pump Tubing, 1/8" ID x 1/4" OD; 25 Ft | Cole-Parmer 50123-55 | This peristaltic silicon tubing withstands repeated clean-in-place cleaning and sterilization and is chemical resistance (*see Figure 1.1.A. Viii.*) |
| **Filter Membrane** |  | The size of the membranes varies in the literature with researchers using 300kDa, 500kDa and up to/combined with 1000kDa molecular weight cut off (MWCO) membrane to isolate EVs (2). The three sources validated in this protocol use 500kDa membrane filter for maximum protein clearance while retaining particles <30nm (figure 3-5). Can be optimized for other MWCO membranes. |
| Repligen MicroKROS 20cm 500kDa PS 0.5mm mL x FLL | Repligen C02-S500-05-N | 500 kDa Repligen MicroKROS membrane (*see Figure 1.2.C*) does not require pre-wetting with alcohol (*see methods 1.1.2 for workflow*) |
| **Cap Assembly** |  | *See figure 1.1.* |
| Cap: 50 mL Conical Centrifuge Tube cap | Sigma CLS430828 | The cap on which the tubing is added with stopcocks. Blunt needles can be adjusted to the tube size for the sample volume (e.g. a 500ml bottle cap if the sample volume is large 100-500ml). |
| 3 × BD Needle Drawing Up Sterile 18G Blunt (1.2 X 38mm) | BD 300204 | The 18G needle hub connects to the stopcocks allowing sample to flow out from the falcon tube. (*see figure 1.1.D*) |
| 2 × BD Microlance 3 Needle Sterile 16G blunt (1.6 x 40mm, white) | BD 300637 | The 16G needle can be inserted into the 18G needle from underneath the cap to enable fixing male luer lock ring with clear tubing (50119-64); then adjusted to the height of the falcon tube for sample to be taken up (*see figure 1.1.G*) |
| 2x Stopcocks: Terumo Stopcock, 3 way, luer lock, sterile, 50/box | Livingstone International TTST21L | Connects tubing and the needle hub (*see figure 1.1.H*). |
| **Connectors:** |  |  |
| 2 × Female Luer Thread Style to 200 Series Barb 1/8in (3.2mm) ID Tubing Animal Free Polypropylene | Chromalytic 0218-PP00-004 | Connected on one end of both long and short arm tube, which connects to the either end of the TFF-membrane (*see figure 1.1.M*). |
| 2 × Male Luer Integral Lock Ring to 200 Series Barb 1/8in (3.2mm) ID Tubing Animal Free Polypropylene | Chromalytic 0418-PP00-004 | Connected on another end of long and short arm tube, which connects to either end of the stopcocks (*see figure 1.1.M*). |
| **Other materials:** |  |  |
| Hot glue gun | Varied; (Anko TY-G1001) | To ensure the system is air-tight for the run and diafiltration (*see figure 1.1.D*). |
| Hammer (or something similar) | Varied | To pin the needles holes in the cap (*see figure 1.1.C*). |
| Tubing clamp for pressure (Hoffman Tubing Clamp) | Varied; Hoffman 30mm stainless steel KL37891 | (*see Figure 1.2.C*) |
| Scissors | Varied | To adjust the length of the tubing. |
| Alcohol wipes | Varied | To clean all external surfaces during set up. |

#### 5.1. Material listed for system components

Individually sourced components, catalogue numbers, and notes for the lab-assembled TFF system are listed in Table 1.

#### 5.2. Step-by-step assembly of the cap

Wipe all external surfaces of the assembly with alcohol wipes throughout.

1. Take a 50ml conical tube cap as the base of the assembly (*see Figure 1.1.B*).
2. Using a hammer (or any heavy object) to pierce three holes through the cap using 18G blunt needles with 1.5cm gap between each hole (*see Figure 1.1.C*).
3. Insert 18G blunt needles in each hole from the top of the cap (*see Figure 1.1.D)*.
4. Apply hot glue around the base of each needle hub, on both the top and underside of the cap, to create an airtight seal capable of withstanding the pressure generated during filtration and diafiltration (*see Figure 1.1.D)* (Note: let the hot glue cool and firm before applying another layer to ensure coverage. When applying hot-glue underside the cap, ensure the edges of the cap are not blocked as the conical tube needs to be closed*). (Caution: Hot glue guns reach temperatures of ∼150-200°C. Molten glue can cause burns on contact with skin and remains hot for several seconds after application. Allow glued needle-cap junctions to cool fully before handling. Use in a well-ventilated area, as heated adhesive can release fumes. Keep the glue gun tip away from tubing and other heat-sensitive components during application)*.
5. Insert a 16G blunt needle into the 18G needle from the tip, from underneath the cap (*see Figure 1.1.G*). Attach a male luer lock to each 16G needle hub connected to 5.2cm of platinum-cured silicone tubing (Cole-Parmer 50119-64) trimmed with scissors to the internal height of the 50ml conical tube (*see Figure 1.1.I & K*). This allows the tube to draw sample from the bottom of the reservoir. *(Caution: Blunt-tip needles (18G, 16G) minimize but do not eliminate puncture risk, particularly during insertion through the cap or when handling loose needles before assembly. Use care when piercing pilot holes with the hammer, as the needle tip is exposed on the underside of the cap during and after insertion. Dispose of any bent, damaged, or contaminated needles in an approved sharps container, not general laboratory waste.)*
6. Attach a 3-way stopcock to each needle hub on top of the cap, providing independent control of the sample inlet, sample outlet, PBS buffer inlet (*see Figure 1.1.J*).
7. Cut two lengths of PharMed BPT peristaltic pump tubing (Cole-Parmer 50123-55) to size, a short arm and a long arm, to connect the cap assembly to the two ports of the hollow fiber membrane (*see Figure 1.1.L*).
8. Fit a female luer barb connector (Chromalytic 0218-PP00-004) to one end of each tube, for attachment to the membrane ports (*see Figure 1.1.M*).
9. Fit a male luer lock ring connector (Chromalytic 0418-PP00-004) to the other end of each tube, for attachment to the stopcocks on the cap (*see Figure 1.1.M*).

#### 5.3. Cleaning and Set-up

1. Clean the external surfaces of the three tubing lines and the needle attached to the cap sequentially with:

- 40 mL Milli-Q water (removes loose dirt/debris)
- 40 mL 70% ethanol (disinfects, removes organic contaminants)
- 40 mL Milli-Q water (rinses ethanol residue)
- 40 mL PBS (equilibrates tubing to physiological buffer conditions before sample contact) (*see Figure 1.2.A & B*)

*Notes: Ensure the reagents are filtered (0.02μm)*.

2. Connect the short-arm and long-arm tubes prepared in the cap assembly (Section 1.2) to the two ports of the 500kDa hollow fiber membrane (Repligen C02-S500-05-N), completing the cap-to-membrane assembly (*see Figure 1.2.C)*.
3. Attach a 50 mL conical tube containing 45 mL PBS to the cap. *Note: this step applies to first-time use of the membrane only. For repeated use, the membrane must first be washed free of the cleaning/storage reagent, for example, if the membrane was stored in 70% ethanol, it should be flushed with filtered Milli-Q water before PBS priming. All reagents introduced to the membrane should be pre-equilibrated to 4°C, matching the sample temperature, before use*.
4. Start the peristaltic pump at 35 mL/min to flush and prime the system. Keep the entire setup inside a 4°C fridge throughout.
5. Back Pressure Set-up: Adjust the Hoffman pressure clamp on the short arm or retentate line to ensure the permeate rate drip is 15-20 drops per 30 seconds (∼75-80ul per drop).
6. Pause the pump when approximately 2 mL of PBS remains in the conical tube, to prevent the membrane drying out and being damaged.

#### 5.4. EV-isolation via TFF

##### Sample preparation

After preprocessing (see Methods 4), all the CM samples (20ml per sample for human MDM and Cerebral Organoid, 7ml for BMDM) were diluted to equal volume of 35 mL with sterile, cold, filtered PBS.

*The following steps are continued from Section 5.3 (after set-up, cleaning and priming):*

7. Remove the PBS tube from the cap and replace with the prepared sample tube (*Figure 1.3.A*).
8. Start the peristaltic pump to begin concentrating the sample. The permeate rate can be expected to drop slightly (more with serum containing media than serum-free media) while concentrating, however, flow rate needs to be maintained within the source-specific target range (Table 2) throughout the run. To maintain the flow rate, incrementally tighten or release the Hoffman clamp to restore the target flow rate and avoid excessive backpressure.
9. Pause the pump once approximately 15 mL of sample remains in the conical tube, to avoid over-concentration and preserve EV integrity.
10. **Diafiltration:** submerge the diafiltration inlet tubing (clear tubing) into a reservoir containing 500 mL PBS. Restart the pump and diafilter with approximately 300 mL PBS over 2-3 hours, to remove free proteins and other soluble contaminants by buffer exchange. Pressure and permeate flow should be monitored to confirm PBS is entering the retentate side dropwise at the same rate permeate is discarded, keeping the retentate volume steady at ∼15 mL throughout; check the flow rate every 15-30 minutes. If PBS does not begin flowing from the buffer inlet, check the tubing is correctly positioned in the bottle; to assist pump priming, gently inject PBS into the inlet line using a syringe, then immediately return the inlet tubing to the PBS bottle.
11. After the 300 mL PBS diafiltration is complete, remove the diafiltration tubing from the PBS reservoir. Restart the pump to concentrate the retained sample down toward 5 mL, then stop the pump.
12. Carefully open the cap and allow all retentate to fall into the collection tube to elute the sample. Final recovered volume should be approximately 7-10mL.
13. **Secondary concentration:** concentrate the eluted EV sample using a 10 kDa MWCO spin filter. In this protocol, technical replicate samples (Human MDM, Organoid) were concentrated to 0.7 mL for use in both TFF-only and TFF+SEC downstream analyses; Mouse BMDM samples were concentrated to 0.5 mL for downstream use on a 70 nm qEV column.
14. Centrifuge at 4000 × g for 20-25 minutes at 4°C, using a pre-chilled centrifuge.

**Table 2:** Target and observed permeate flow rates.

| Source | Average flow rate (drops/30seconds) | Flux, J (LMH), see equation (1) below | Notes: |
| --- | --- | --- | --- |
| PBS priming (for all sources, pre-use QC check) | 15-20 | 48-68 | Confirms membrane is free of residual fouling from prior use before sample is introduced. If priming flow rate falls below 14-15 drops/30 sec, the membrane is considered fouled beyond adequate cleaning and is retired from use (see PBS Flux Recovery, Table 3). |
| Human MDM | 10-11 | 32-37.4 | Hoffman clamp required incremental tightening throughout the run to maintain target rate as resistance developed. |
| Human cerebral organoid | 15-16 | 48-54.4 | Minimal clamp adjustment required. |
| Mouse BMDM | 11-12 | 35-40.8 | Minimal to moderate clamp adjustment required. |

*General notes: CM and CM-EVs were always kept cold and after isolation, were kept in ice for aliquoting prior to storage at -80°C. All EV aliquots were stored in protein lo-bind Eppendorf tubes*.

#### 5.5. Post-sample cleaning

1. Remove the sample tube and replace with a conical tube containing 45 mL filtered Milli-Q water. Start the pump to rinse the system, removing residual buffer and sample components.
2. Replace the water tube with a conical tube containing 45 mL 0.5N NaOH. Adjust the Hoffman clamp to maintain a backpressure corresponding to 15-20 drops/30 sec. *(Optional: Stop the pump once the NaOH has passed through the system, then leave the system static with NaOH inside for a 1-hour incubation, or overnight if running the sample the next day.* NaOH denatures residual protein, disinfects the system, and removes stubborn biological residues.)
3. Rinse the system with 45 mL Milli-Q water to remove NaOH residues, then either rinse with 45 mL 70% ethanol and fill the membrane with 70% ethanol for storage at 4°C, or prime with PBS to start the next run of the sample. *(Note: If the membrane is to be stored in ethanol, ensure the membrane lid is fixed on both ends and keep at 4°C. The membrane can be reused up to 4-5 times.)*
4. The disassembled tubing needs to be washed twice with 50 mL Milli-Q water (to remove residual cleaning agents), and once with 50 mL 70% ethanol before storing at room temperature. *(Note: The tubing needs to be cleaned sequentially as stated in Section 1.3 before every use and should be changed after 6-10 samples, and after checking for its integrity, e.g. loose stopcock, leakage etc.)*

#### 5.6. Transmembrane pressure monitoring via permeate flow rate

This protocol does not have a pressure gauge added to the set up (which, can in theory be added in the design). Here, transmembrane pressure (TMP) was inferred by monitoring permeate flow rate. Permeate droplet volume was calibrated at ∼75-80 µL per drop by measuring volume of droplets at priming, sample concentration and diafiltration steps. Permeate flow rate was monitored throughout each run by counting drops over a 30-second interval every 15-30 minutes. Target flow rates were maintained via adjustment of the Hoffman clamp on the retentate line: 15-20 drops/30 sec during PBS priming, source-specific ranges during sample processing (Table 2), and 15-20 drops/30 sec during cleaning, corresponding to fluxes shown in Table 2.

Prior to each use, membranes underwent a PBS priming step to confirm system integrity (Method 5.3). A priming flow rate below 15 drops/30 sec (corresponding to a PBS Flux Recovery, PFR, of approximately 75% relative to the pristine membrane baseline) was used as the criterion for membrane retirement, indicating fouling not resolved by the standard cleaning cycle (Method 5.5). Once a sample was introduced, permeate flow rate dropped and stabilized at source-dependent levels (Table 2).

#### 5.7. Membrane flux rate and PBS Flux Recovery (PFR)

Permeate flux (J), the rate at which liquid passes through the membrane per unit area, is calculated as:

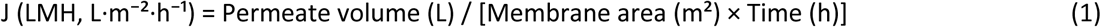

For operational monitoring via drop counting per 30 seconds, using a calibrated droplet volume, which we measured 75-80 µL per drop and membrane surface area of 28 cm² (0.0028 m²), this simplifies to:

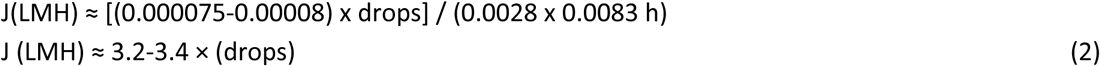

We calibrate this against PBS permeate flux for each repeat use of the membrane, using PBS Flux Recovery (PFR) (Table 3) rather than the conventional Water Flux Recovery (WFR), since PBS, not water, is the sole buffer used throughout system priming, sample processing, and cleaning in this protocol:

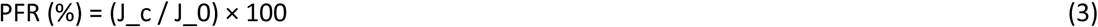

where J_c is the PBS permeate flux measured after cleaning, and J_0 is the PBS permeate flux of the pristine, first-use membrane.

**Table 3:** Expected flux and PBS Flux Recovery (PFR) and cleaning values by membrane condition.

| Stage | Flow rate (drops/30 second) | Volume | Flux (LMH) | PFR (%) |
| --- | --- | --- | --- | --- |
| Acceptable priming range | 15-20 | 75-80ul | ~48-68 | 75-100 |
| Acceptable cleaning range* | 12-20 | 75-80ul | ~38-68 | 60-100 |

\* Droplet volume calibrated at 75–80 μL throughout. PFR calculated relative to each membrane's pristine, first-use PBS flux ( $J_0$ ). A pristine membrane typically primes at 15-20 drops/30 sec, corresponding to that membrane's 100% baseline. Flow rate during cleaning can start low, reflecting residual fouling immediately after the sample run, and is expected to recover gradually across the wash sequence depending on sample type; it is not used as a retirement criterion.

### 6. Size Exclusion Chromatography (IZON 70m qEV column)

Following TFF concentration, TFF-EVs were further processed by SEC using an IZON 70 nm qEV column to remove co-concentrated non-EV proteins (serum-containing preparations) or to further refine the EV population (serum-free preparations).

500 µl of TFF-EVs were thawed on ice. The SEC column was flushed with 17 mL of PBS and then loaded with 500 µl of TFF-EV. Next, 2 mL of PBS was added and collected within the same Eppendorf tube as buffer volume. Then, 400 µl of PBS was added to elute each fraction, from fraction 1 to 6.

Fractions 1, 2 and 3 were pooled together as EV-enriched fractions (Supplementary Figure 1–2).

The SEC column was reused up to 5 times prior to discarding. Between uses, the column was cleaned according to the manufacturer’s instructions and stored at 4°C in an upright position.

### 7. Transmission Electron Microscopy (T12, Sydney Microscopy and Microanalysis)

EVs were stained with 2% Uranyl Acetate on a 300-mesh Copper grids (GSCU300CC-50) with respective dilution (see Supplementary Table 1). The grids were imaged on Tecan T12 transmission electron microscope at the Sydney Microscopy and Microanalysis facility.

### 8. Nanoparticle Tracking Analysis (Zetaview)

Particle size distribution and concentration of EV preparations were determined by nanoparticle tracking analysis (NTA) using a ZetaView PMX-120 (Particle Metrix, Germany) at Sydney Analytical equipped with a 488 nm laser operating in scatter mode. All measurements were performed at 25°C with pH 7.0 and sensitivity set to 80 with shutter 100.

#### Sample preparation for NTA

All diluent solutions were prepared using phosphate-buffered saline (PBS) filtered through a 0.02 µm syringe filter (Whatman, UK) immediately prior to use to eliminate background particles. EV preparations were diluted in 0.02 µm-filtered PBS to achieve a particle concentration within the optimal detection range of the instrument (approximately 100-400 particles per frame) (see Supplementary Table 3).

#### Instrument acquisition settings

Each measurement consisted of one cycle across 11 positions. Particle tracking was performed with the following analysis parameters: minimum area 10, maximum area 1000, minimum brightness 20, size class interval 10 nm. Positions with deviant scatter patterns were automatically removed by the Zetaview software. A minimum of 200 traced particles per measurement was targeted, measurements falling below this threshold are noted (see Supplementary Table 2). Standard acquisition was captured with scatter filter (sensitivity 80, shutter 100). Fluorescent acquisition was captured with 500nm filter wavelength (sensitivity 95, shutter 100).

#### Blank measurements

Instrument background was assessed by measuring 0.02 µm-filtered PBS (instrument blank) between each sample throughout the measurement session to confirm the absence of background particle contamination. All blank measurements were below the instrument detection threshold prior to each sample acquisition.

#### Cell Mask Green (CMG) membrane staining

To assess the proportion of membrane-enclosed particles, a subset of TFF+SEC preparations (human macrophage n=3; cerebral organoid n=2) were stained with Cell Mask Green (CMG) plasma membrane stain (Thermo Fisher Scientific, C37608). CMG stock solution was diluted 1:200 in 0.02 µm-filtered PBS. For staining, 4.5 µl of EV preparation was combined with 0.5 µl of 1:200 CMG, incubated for 1 h RT, dark and topped to 500 µl with 0.02 µm-filtered PBS (final dilution 1:100; final CMG concentration 1:100,000 of stock). After incubation with CMG, all the EVs were kept on ice and captured by NTA within 1-2 h of staining. CMG staining was not performed on TFF-only preparations due to the high background signal from co-concentrated serum proteins interfering with fluorescence detection, nor on mouse BMDM preparations.

#### Data analysis and concentration calculation

Raw NTA output files were exported from ZetaView software (version 8.05.16 SP3) as size distribution tables containing particle concentration (particles/cm³) per 5 nm size bin across a measurement range of 2.5-1000 nm. Dilution-corrected concentration values were used for all subsequent analyses. Total particle concentration in the EV stock preparation (particles/mL) was calculated by integrating the size distribution across all size bins:

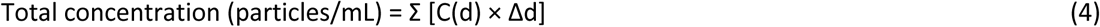

where C(d) is the dilution-corrected concentration at size bin d (particles/mL) and Δd is the bin width (5 nm).

Total particles in each EV preparation were calculated by multiplying the total concentration by the final EV resuspension volume and normalized to the volume of conditioned medium processed (particles per mL of conditioned medium). Particle-to-protein ratios (particles/µg) were calculated using the total protein content (µg) determined by Qubit protein assay quantification (see Supplementary Table 1).

#### Reporting

NTA measurements are reported in accordance with MISEV2023 guidelines (1). Mode size, D10, D50 (median), and D90 values are reported per replicate alongside total particle concentration and particle-to-protein ratio. Individual size distribution traces are shown for each replicate. Summary statistics (mean ± SD) are reported for conditions with three replicates, whereas individual values are reported without summary statistics for conditions with two replicates (see Supplementary Table 2).

### 9. Protein Quantification by Qubit

A Qubit Protein Assay (Thermo Fisher Scientific Australia Pty Ltd, Q33212) was used for protein quantification of protein concentration, as per manufacturer’s instructions.

### 10. Mass Spectrometry

EV peptides from human MDM and Human Cerebral Organoid were processed in the same batch during protein digest and data acquisition, and Mouse BMDM and BMDM_LPS processed together in another batch.

#### 10.1. EV proteome for LC-MS/MS

The EV samples (4.5-49 µg; see Supplementary table 1) were resuspended in 0.2% (*w/v*) RapiGest SF^TM^ (25uL; Waters, 186001861), followed by 0.05 mol/L of TEAB (Sigma, T7480) to a total volume of 500µl (Figure 4, step 2). To aid EV lysis and protein resuspension, the EV samples were incubated at 95°C for 5 minutes and sonicated twice using a step-tip probe at 20% amplitude for 15 seconds (EVs were rested on ice between pulses). The proteins were reduced in 12 mM tris(2-carboxyethyl) phosphine (TCEP, (C4706-2G, Sigma-Aldrich) at 60°C for 30 minutes. Alkylation was performed with 50 mM Iodoacetamide (Sigma, A3221) in the dark at RT for 30 minutes (Figure 4, step 4). Sample pH was adjusted to 7.5 using 0.05 M of TEAB, and sequencing-grade trypsin was added to the sample at a ratio of 1:30 (w/w) trypsin: protein ratio (Promega, V511A) and incubated overnight at 37°C.

After 16 hours, Rapigest was cleaved by acidification of the sample with 50% (v/v) formic acid (FA) to pH to <2, followed by incubation at 37°C for 30 minutes. The rapigest was clarified from the sample by centrifugation twice (13,000 rpm, 10 minutes, 4°C). EV peptides were desalted using a 1cc Hydrophilic-lipophilic balance (HLB) column (Waters,186000383; see Figure 4, step 6). The EV peptides were resuspended with 1mL of 0.1% (v/v) FA. After conditioning the HLB column with 1mL of 100% acetonitrile (ACN), 1 ml of 50%ACN/0.1%FA and 1 mL of 0.1%FA, the EV peptides were passed through the HLB column 5 times to ensure full retention of the peptides on the HLB sorbent. The HLB column was then washed three times with 1mL of 0.1% (v/v) FA, and peptides were eluted into a protein lobind tube by 500μL of 50% ACN/0.1% FA, followed by 200μL of 70%ACN/0.1% FA. A final Qubit quantification was performed on the desalted peptides, followed by drying for 9 h using a centrifuge evaporator at 45°C.

#### 10.2. Liquid Chromatography-Tandem Mass Spectrometry (LC-TMS)

The dried EV peptide pellet was reconstituted to a final concentration of 0.1 μg/μL with 3% acetonitrile (Sigma, 34851) and 0.1% FA (Thermo, 85178), sonicated in a water bath for 10 min and centrifuged at 10,000 × g for 10 min at 4 °C. Peptides (3 μl; 300 ng) were separated on an in-house packed column (75 μm × 450 mm, 1.9 μm ReproSil Pur 120 C18, Dr. Maisch GmbH, Germany) using a Vanquish Neo liquid chromatograph system (Thermo Fisher Scientific) coupled to a Thermo Q Exactive HF-X mass spectrometer operated in positive nano-electrospray ionisation mode. Peptides were separated at a flow rate of 0.3 μL/min with a gradient of 0.1% FA in H2O (Solvent A) and 0.1% FA in 80% ACN (Solvent B), linearly increasing from 5% to 35% B over 66 min, then to 60% B over 2 min and 98% B over a further 2 min, followed by a column wash at 98% B (flow rate 0.5 μL/min) for 5 min prior to re-equilibration. Data were acquired (Figure 3) in data-independent acquisition (DIA) mode. MS1 spectra were acquired at a resolution of 60,000 across a detectable mass-to-charge (m/z) range of 350–1650, and MS2 spectra were acquired at a resolution of 30,000 using 20 isolation windows across the 75- minute gradient. Proteomic data were extracted using Xcalibur data acquisition software (Thermo Fisher Scientific).

##### DIA analysis

Thermo RAW data files were analysed in DIA-NN (version 2.2.0) software against Uniprot FASTA databases (either *Homo sapiens* or *Mus musculus* (Swiss-Prot reviewed uniprot database 2026_03_18)(16). Predicted spectral libraries were generated for the two species using the raw DIA files, and then DIA data from the raw files were extracted against the respective predicted spectral library with the reannotation function enabled. Data were searched across the precursor range of 350-1650 m/z and a fragment ion range of 300-2000 m/z, with trypsin missed cleavage of 2.

### 11. Data Analysis and statistics

All proteomic and particle data analysis was performed in R. Protein group intensities (DIA-NN pg_matrix output) were log2-transformed prior to all downstream analyses; no imputation was performed.

The number of proteins detected per replicate was compared between TFF-only and TFF+SEC. Reproducibility was assessed by the coefficient of variation (CV%, SD/mean × 100) of raw protein intensities, calculated for proteins detected in all replicates of a condition and shown as violin plots with median CV% reported per condition; CV was not calculated for organoid EVs (n=2). Principal component analysis was performed using *prcomp() (scale. = TRUE, center = TRUE)* in R (version 2026.01.1+403), on proteins detected in all replicates being compared, with percentage variance explained by PC1 and PC2 reported. Particle-to-protein ratio (total particle number by NTA / total protein content by Qubit) was used as the purity metric and is shown as mean ± SD with individual replicate values; the human MDM comparison (Figure 3M) was tested by paired t-test (n=3, df=2).

A heatmap of log2 protein intensities was generated using *ComplexHeatmap* and *circlize*, with markers grouped into the five MISEV2023 Table 3 categories (1). EV identity was benchmarked against the Vesiclepedia top 100 marker list (17), visualized as Venn diagrams using *ggVennDiagram*.

Differential protein abundance between TFF-only and TFF+SEC (human macrophage, organoid) was tested using limma on intensities normalized with variance-stabilizing normalization (vsn); values are reported as fold change (glog) rather than log2 fold change, reflecting the generalized-log transformed used. Proteins were significant at |fold change (glog)| > 1 and adjusted p < 0.05 (Benjamini-Hochberg). Gene Ontology enrichment of significant proteins was performed using gprofiler2 (GO:BP, GO:CC, GO:MF; g:SCS correction; whole-genome background), excluding the low-specificity term “protein binding” and collapsing near-synonymous terms sharing more than 80% gene overlap; the top two enriched terms per category are reported.

Software: dplyr, tidyr, ggplot2, stringr, patchwork, scales, ComplexHeatmap, circlize, ggVennDiagram, vsn (v3.78.1), limma (v3.66.0), gprofiler2 (v0.2.4). NTA data were visualised in GraphPad Prism.

### 12. EV-TRACK

We have submitted all relevant data of our experiments to the EV-TRACK knowledgebase (EV-TRACK ID: EV260077) (Van Deun J, et al. *EV-TRACK: transparent reporting and centralizing knowledge in extracellular vesicle research.* Nature methods. 2017;14(3):228-32).

## Results

To validate the lab-assembled TFF system (Figure 1), we tested it against three different sources of CM: 1) human MDM cultured with EVdep-HuS; 2) human cerebral organoids cultured without serum; and 3) mouse BMDM, unstimulated and LPS-stimulated, cultured with EVdep-FBS. All samples underwent a standardised pre-processing, isolation and analysis workflow (Figure 2), allowing assessment of TFF system performance across serum-containing and serum-free media, and across species. Technical replicates of human MDM (n=3, Figure 3) and human cerebral organoid (n=2, Figure 4), allowed assessment of TFF-system reliability and EV isolation reproducibility. Biological replicates of mouse BMDM cultured under unstimulated (n=3) and LPS-stimulated (n=3) conditions, enabled the evaluation of whether the TFF workflow could capture biologically meaningful differences in EV profiles between two distinct biological conditions (Figure 6), demonstrating the suitability of our lab-assembled TFF systems for comparative EV proteomic studies.

**Figure 1:**
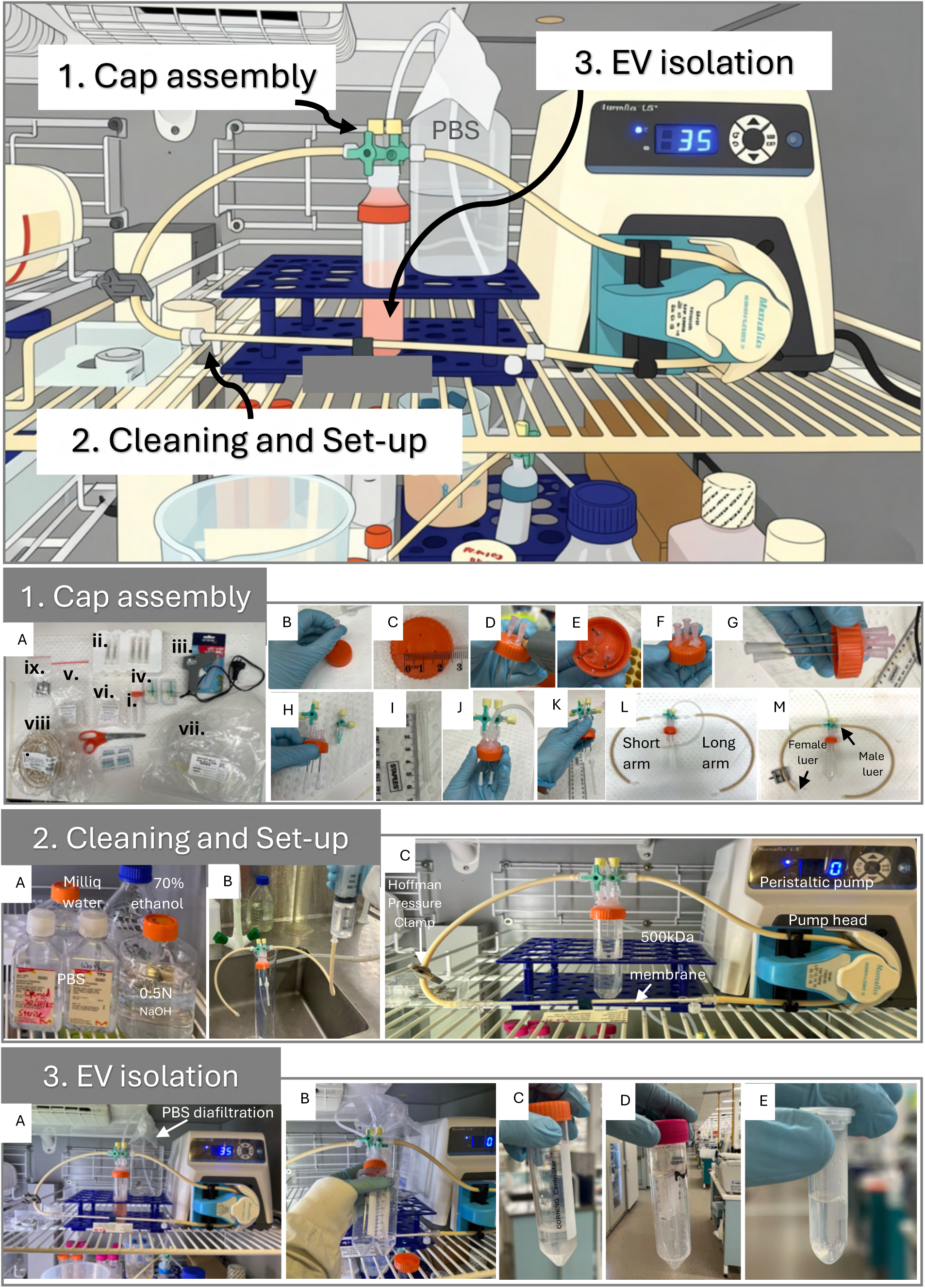
Lab-assembled TriPort-TFF system construction and operation. Top: Overview of the assembled TFF system showing the three operational stages. (1) Cap assembly: components and sequential assembly of the TFF cartridge cap with three inlet/outlet ports. (1.A) Required components including luer-lock fittings and tubing for TFF cartridge. (1.B-M) Sequential cap assembly steps with tubing connections. (2) Cleaning and set-up: system priming and equilibration. (2.A) Cleaning reagents include milliq water, 70% ethanol, 0.5N sodium hydroxide and Phosphate buffer saline (PBS) (all 0.02μm filtered). (2.B-C) System washing and apparatus assembled system during set-up. (3) EV isolation: shows TFF concentration of conditioned media. (3.A-E) Sequential steps during EV isolation including pump operation, sample loading, concentration, and retentate collection. All procedures were performed at 4°C. See methods and Table 1 for detailed step-by-step protocol.

**Figure 2.**
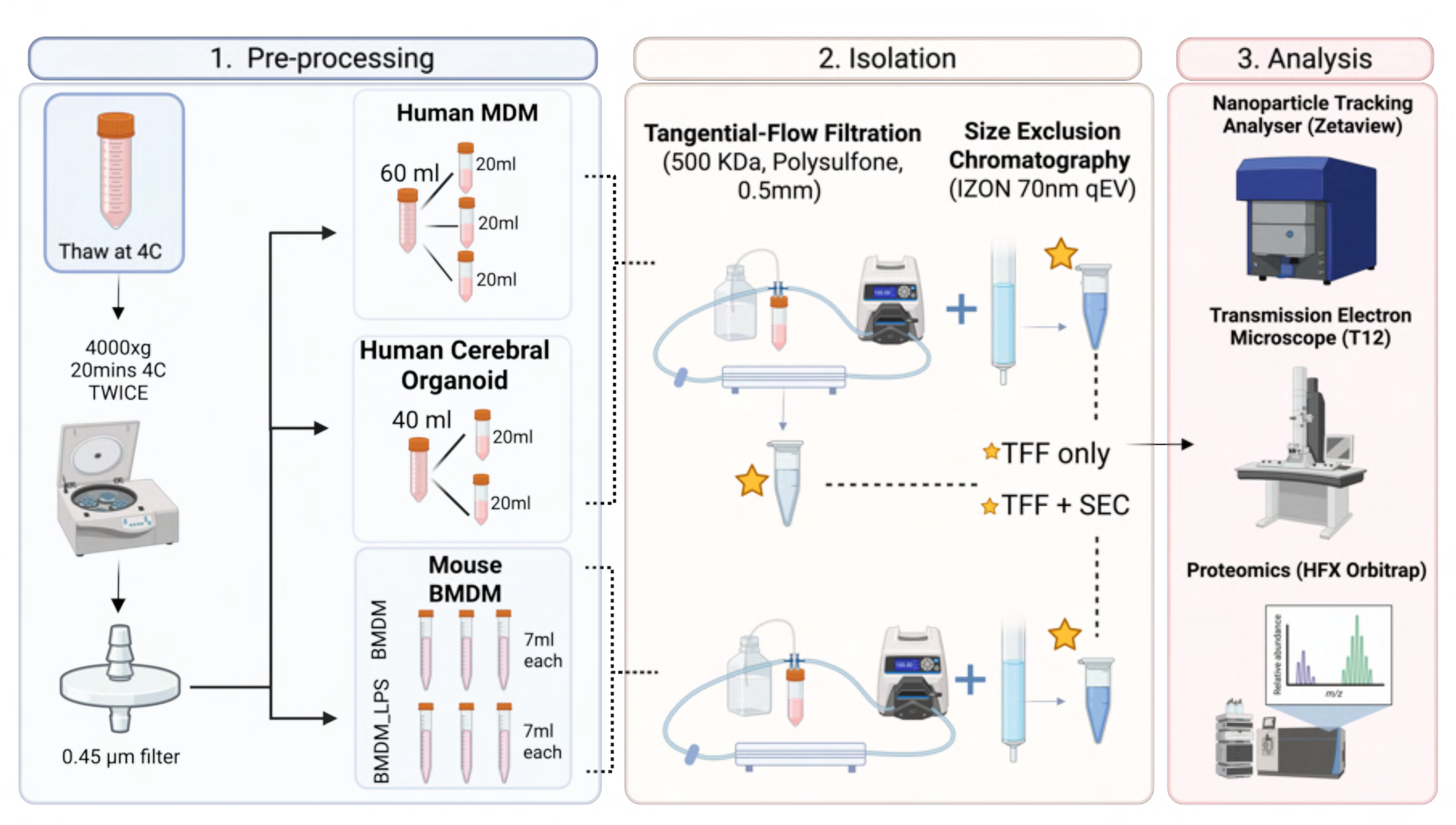
Experimental workflow for EV-isolation from diverse cell culture systems using TriPort-TFF and SEC. Conditioned media (CM) from three biologically distinct cell culture sources was processed through a standardized pre-processing and isolation pipeline. (1) Pre-processing: Frozen CM was thawed at 4°C overnight and subjected to sequential low-speed centrifugation (4,000 × g, 20 min, 4°C, twice) to remove cellular debris, followed by filtration through a 0.45 µm syringe filter. (2) Isolation: Clarified CM was processed using a lab-assembled tangential flow filtration (TFF) system equipped with a 500 kDa MWCO polysulfone hollow-fibre membrane. Human MDM CM (60 mL, split into 3 × 20 mL technical replicates) and cerebral organoid CM (40 mL, split into 2 × 20 mL technical replicates) were processed by TFF alone (TFF-only) and with subsequent size exclusion chromatography (TFF+SEC) using an IZON 70 nm qEV column. Mouse BMDM CM (7 mL per biological replicate; BMDM n=3, BMDM_LPS n=3) was processed by TFF+SEC only. Gold stars indicate aliquots collected for downstream analysis. (3) Analysis: EV preparations were characterized by nanoparticle tracking analysis (NTA) (ZetaView, Particle Metrix), transmission electron microscopy (TEM) (Tecnai T12), and data-independent acquisition mass spectrometry-based proteomics (HF-X Orbitrap).

**Figure 3:**
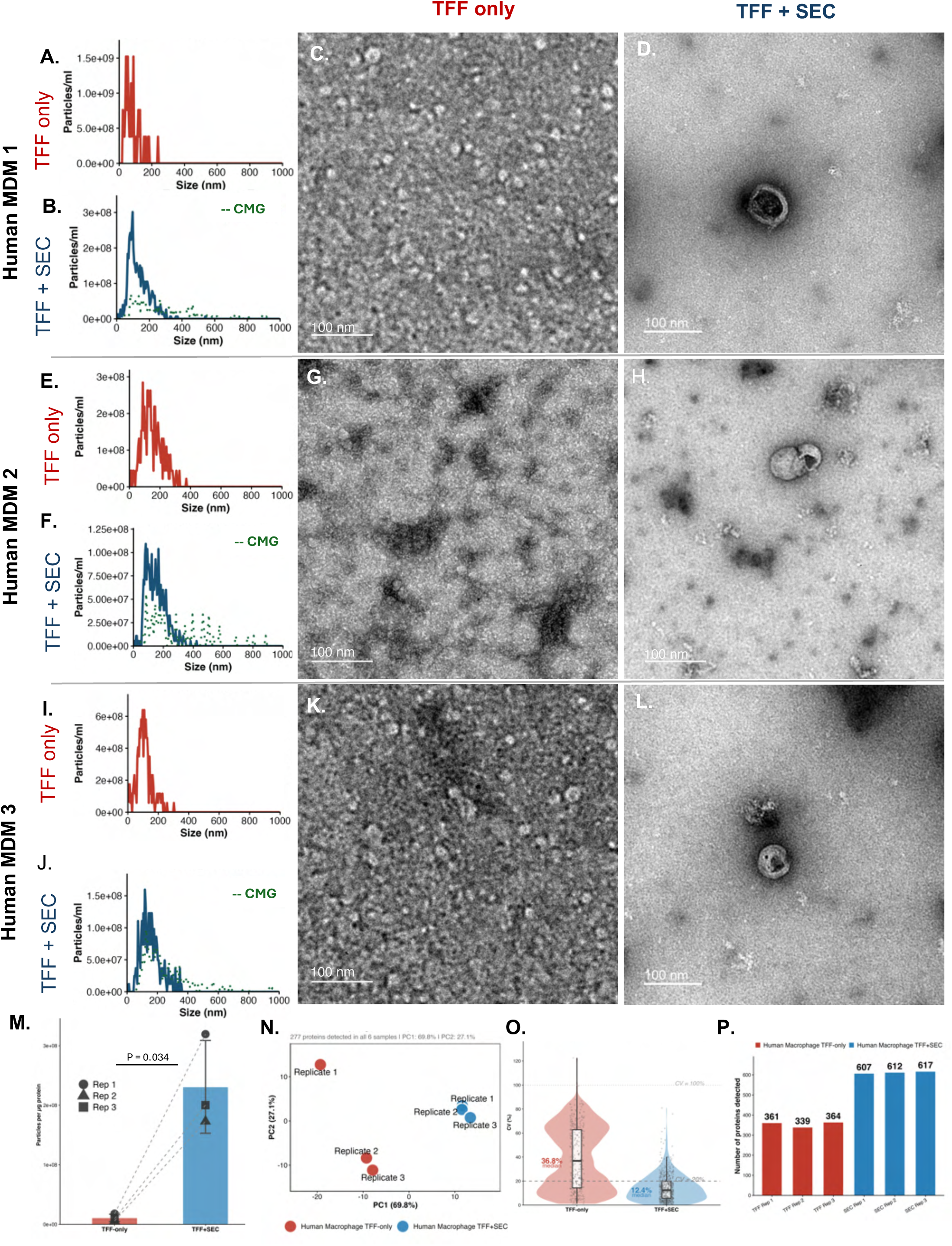
Enrichment of extracellular vesicles from serum-supplemented human MDM conditioned media by TriPort-TFF combined with SEC. NTA and TEM results of Human MDM CM replicate 1 (A-D), replicate 2 (E-H), replicate 3 (I-L) pooled from day 3 and day 7, and split into equal volumes of 20ml, are shown. (A, E, I) NTA particle size distributions for TFF-only preparations (red) of independent technical replicates (n=3). (B, F, J) NTA particle size distributions for TFF+SEC preparations (blue) is shown with membrane-dye (CMG 1:200)-positive particle (light blue dotted line) indicating the proportion of intact membrane-bound particles. (C, G, K) Representative TEM micrographs of TFF-only preparations. Scale bar = 100 nm. (D, H, L) Representative TEM micrographs of TFF+SEC preparations, showing vesicles with cup-shaped morphology. Scale bar = 100 nm. (M) Particle-to-protein ratio (total particles/µg protein) for TFF-only versus TFF+SEC preparations across n=3 independent replicates, demonstrating increased purity following SEC (paired t-test, t = -5.31, df = 2, p = 0.034). Data shown as mean ± SD; individual replicate values connected by dashed lines. (N) Principal component analysis (PCA) of log-transformed LC-MS/MS protein intensities, showing separation between TFF-only (red) and TFF+SEC (blue) preparations across replicates. (O) Coefficient of variation (CV%) of protein intensities across replicates for TFF-only versus TFF+SEC preparations, shown as violin plots with embedded boxplots. Median CV = 36.8% (TFF-only) versus 12.4% (TFF+SEC), indicating improved quantitative reproducibility following SEC. (P) Number of proteins identified per replicate for TFF-only (red) versus TFF+SEC (blue) preparations.

**Figure 4:**
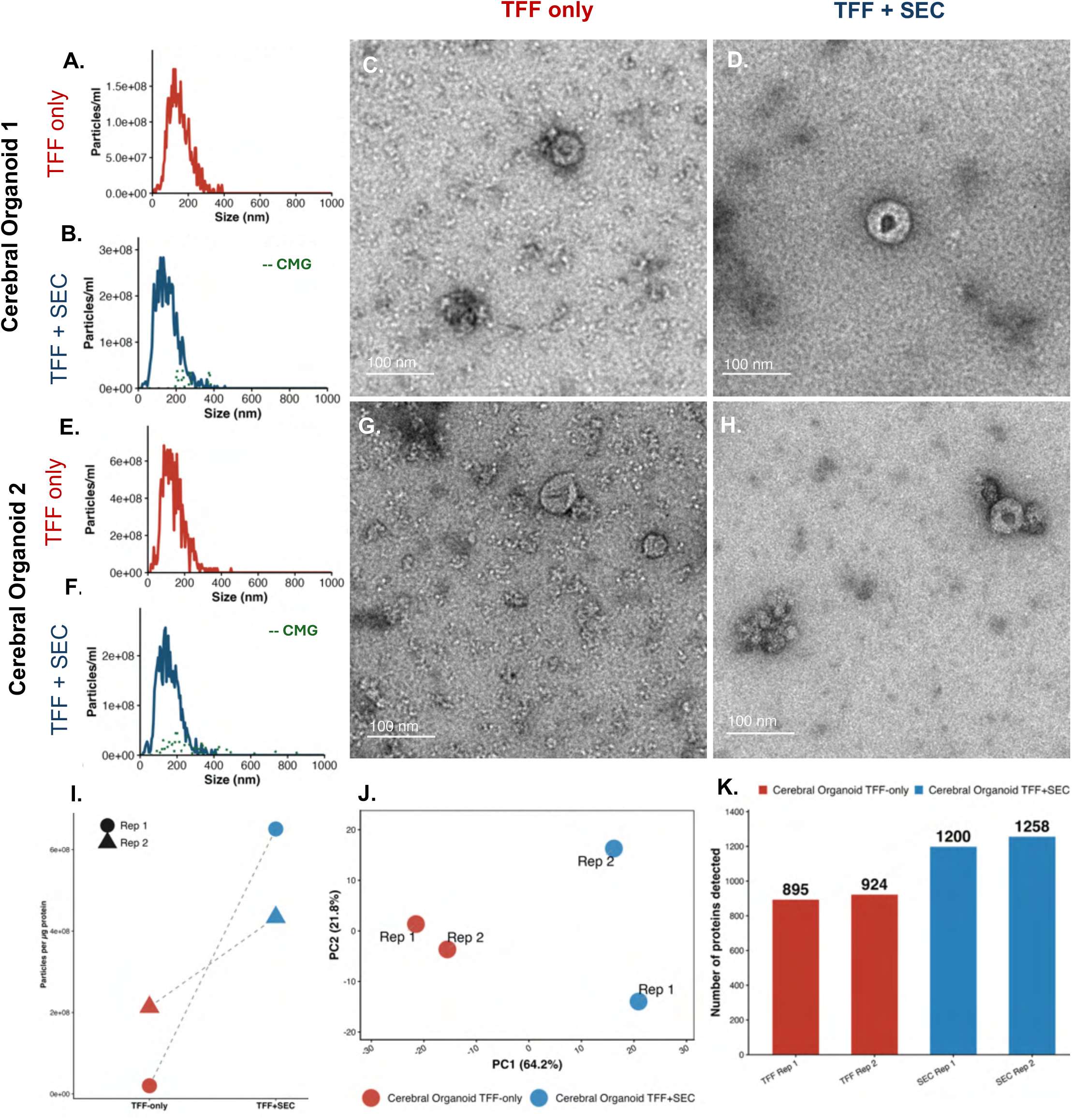
Isolation of extracellular vesicles from serum-free human cerebral organoid conditioned media by TriPort-TFF and altered protein profile following SEC. NTA and TEM results of replicate 1 (A-D), replicate 2 (E-H) from human cerebral organoid CM pooled from day 90-100, and split into equal volumes of 20ml, are shown. (A, E) NTA particle size distributions for TFF-only preparations (red) of independent technical replicates (n=2). (B, F) NTA particle size distributions for TFF+SEC preparations (blue), of independent technical replicates (n=2) is shown with membrane-dye cell-mask green (CMG 1:200)-positive particle (light blue dotted line) indicating the proportion of intact membrane-bound particles. (C-G) Representative TEM micrographs of TFF-only preparations. Scale bar = 100 nm. (D, H) Representative TEM micrographs of TFF+SEC preparations, showing vesicles with cup-shaped. Scale bar = 100 nm. (I) Particle-to-protein ratio (total particles/µg protein) for TFF-only versus TFF+SEC preparations across the replicates, demonstrating increased purity following SEC post TFF. (J) Principal component analysis (PCA) of log-transformed LC-MS/MS protein intensities, showing separation between TFF-only (red) and TFF+SEC (blue) preparations across replicates. (K) Number of proteins identified per replicate for TFF-only (red) versus TFF+SEC (blue) preparations.

**Figure 5.**
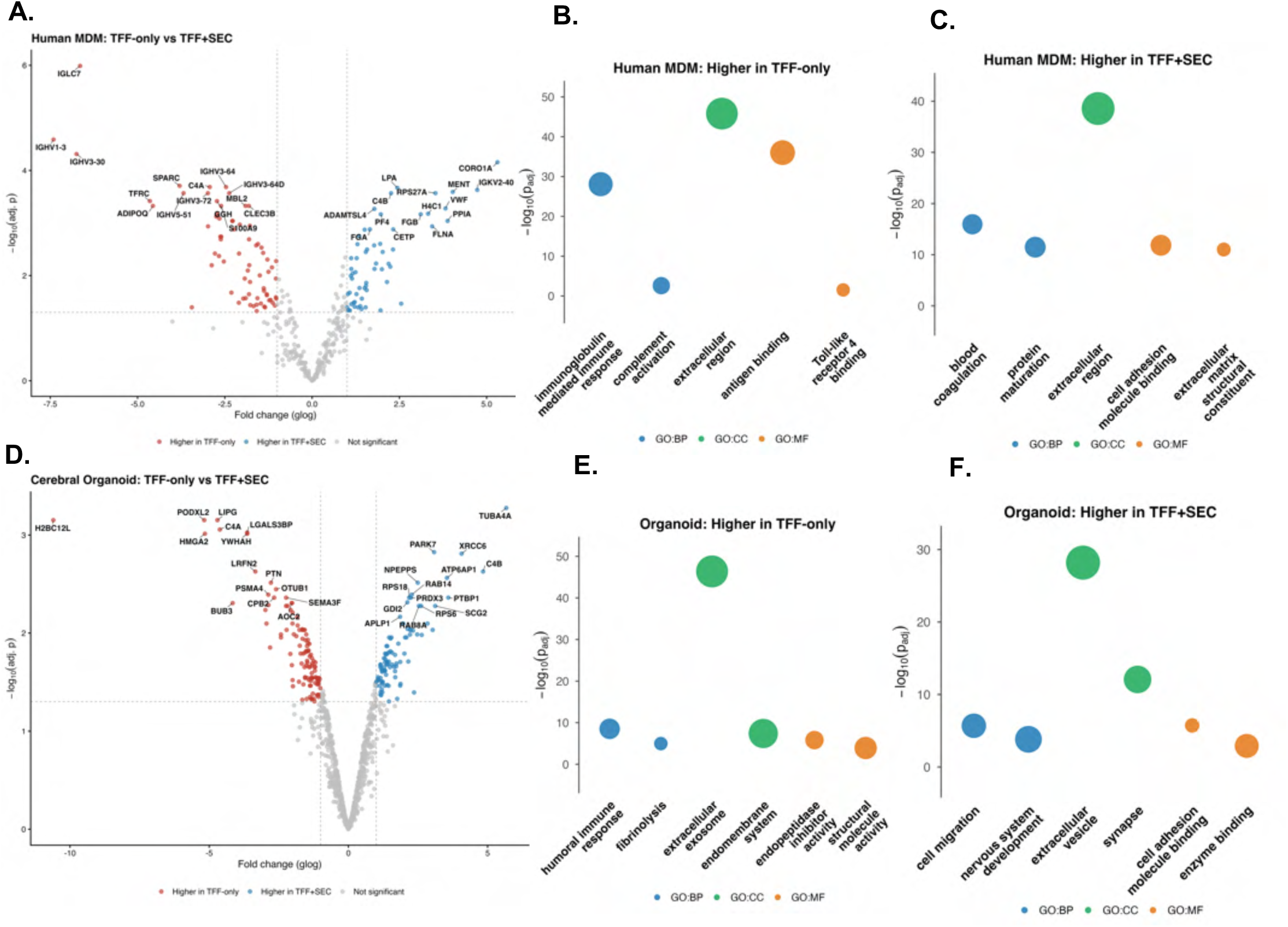
Differential protein abundance and functional enrichment following SEC post-TFF in CM from Human MDM and human cerebral organoid EV preparations. (A, D) Volcano plots of differential protein abundance between TFF-only and TFF+SEC preparations for (A) Human MDM (n=3 technical replicates per condition; 71 significantly higher in TFF-only, 60 significantly higher in TFF+SEC) and (D) human cerebral organoid (n=2 technical replicates per condition; 116 significantly higher in TFF-only, 93 significantly higher in TFF+SEC). Dashed lines indicate the thresholds used to define differential abundance (|fold change (glog)| > 1, adj. p < 0.05). Proteins significantly higher in TFF-only preparations are shown in red, and higher in TFF+SEC in blue, non-significant in grey. The 15 most significant proteins per direction (ranked by adjusted p-value) are labelled with their gene symbol. (B, C, E, F) Gene Ontology enrichment of the differentially abundant proteins from the corresponding volcano plot: higher in TFF-only (B, E) and higher in TFF+SEC (C, F), for Human MDM (B, C) and Cerebral Organoid (E, F). Terms are shown for biological process (GO:BP, blue), cellular component (GO:CC, green), and molecular function (GO:MF, orange); dot size reflects the number of proteins overlapping each term, and the y-axis shows -log10 (adjusted p-value).

**Figure 6:**
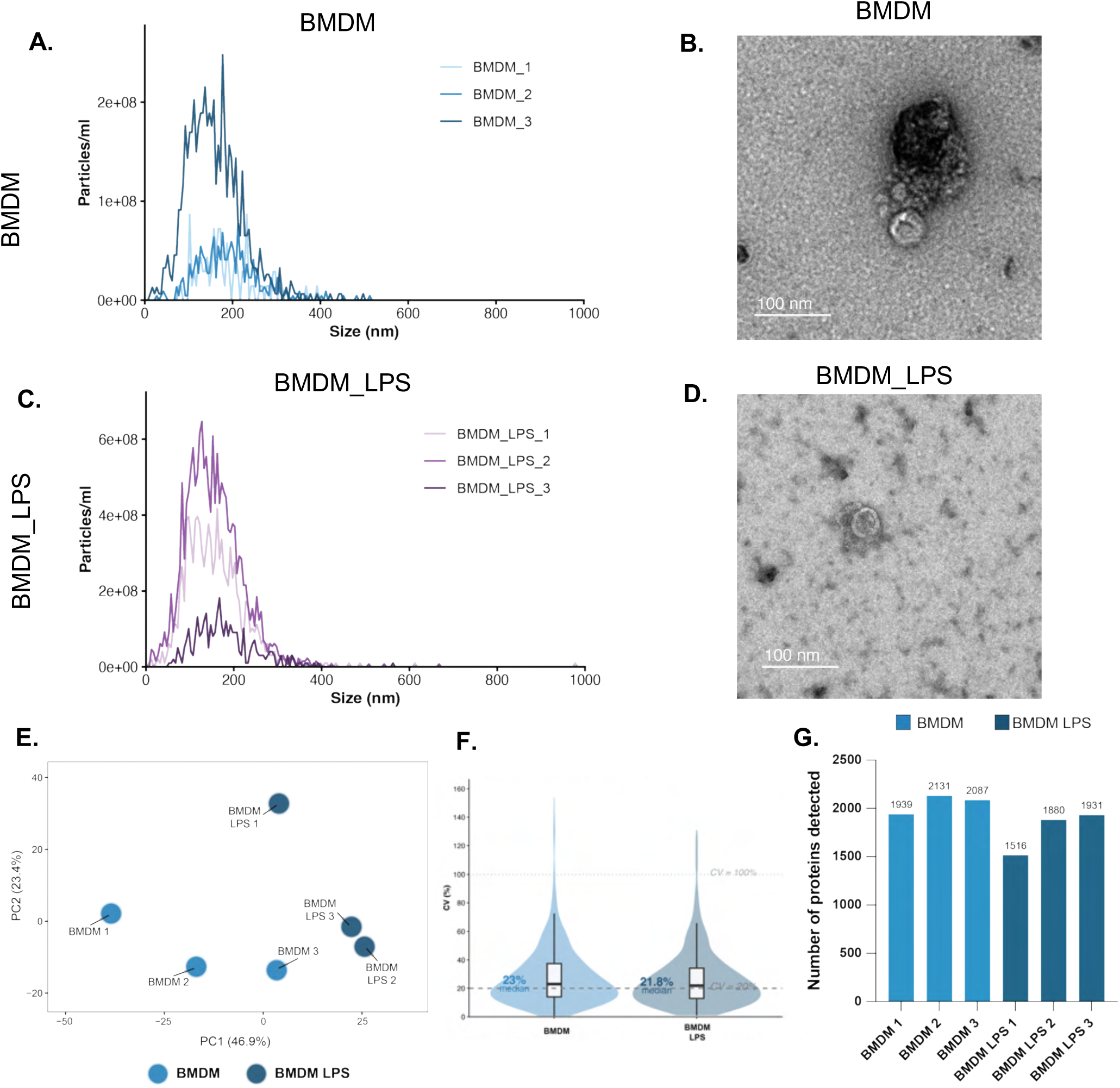
Isolation of extracellular vesicles by TriPort-TFF with SEC from mouse BMDM conditioned media with and without LPS stimulation. (A) NTA particle size distributions for BMDM, three independent biological replicates (light to dark blue). (B) Representative transmission electron micrograph of BMDM-EVs. Scale bar = 100 nm. (C) NTA particle size distributions for mouse BMDM stimulated with LPS, BMDM_LPS, three independent biological replicates (light to dark purple). (D) Representative transmission electron micrograph of BMDM_LPS EVs. Scale bar = 100 nm. (E) Principal component analysis (PCA) of log-transformed LC-MS/MS protein intensities (PC1 = 46.9%, PC2 = 23.4%), showing separation between BMDM and BMDM_LPS across biological replicates from independent animals. (F) Coefficient of variation (CV%) of protein intensities across biological replicates for BMDM versus BMDM_LPS, shown as violin plots with embedded boxplots. (G) Number of proteins detected per sample for BMDM and BMDM_LPS. Biological replicates derived from independent animals, n = 3 per condition.

### Lab-assembled TFF system isolates EVs from serum-containing human MDM conditioned media but co-concentrates abundant serum protein

For human MDM-derived EVs, particle analysis using NTA showed considerable variation in particle yield between replicates: TFF-only EV preparations ranged from 2.22×10^10^ to 5.59×10^10^ total particles, with broad size distributions spanning approximately 50 to 300 nm (Figure 3A, E, I, Supplementary Table 2). In contrast, TFF+SEC EV preparations had lower counts ranging from 5.20×10^9^ to 1.03×10^10^ particles, with a sharper size profile dominated by a single peak spanning 75 and 150 nm (Figure 3B, F, J, Supplementary Table 2). Further, TFF-only human MDM EVs preparations had higher protein concentrations (4500-5788 µg/ml) compared to TFF+SEC (75.2-80.4 µg/ml) (Supplementary Table 1), indicating probable co-concentration of serum-abundant proteins alongside EVs during TFF.

Evidence for SEC removal of non-vesicular co-isolates from TFF preparations is supported by CMG membrane-staining which showed that 35% ± 12% of particles in TFF+SEC preparations were membrane-enclosed. CMG staining was negative on the corresponding TFF-only Human MDM derived EV sample using the same 1:200 dye dilution (Supplementary Table 5). TEM further supported these findings, with the TFF-only preparations exhibiting dense, heterogeneous material adsorbed to the microscopy grids, consistent with substantial co-concentration of non-vesicular protein material in the EV preparation (Figure 3C, G, K). By contrast, for TFF+SEC EV preparations, microscopy grids showed adsorbed EVs with characteristic cup-shaped morphology and there was a lower abundance of protein contaminants (Figure 3D, H, L).

To further characterize EV enrichment following SEC, the TFF-only retentate was compared against individual SEC elution fractions (F1-F4) by NTA and proteomics (Supplementary Table 5, Supplementary Figures 1-2). The particle-to-protein ratio, calculated as a metric of EV purity, increased significantly (paired t-test, t=−5.31, df=2, p=0.034) from an average of 1.09×10^7^ particles/µg in TFF-only preparations to 2.31×10^8^ particles/µg in TFF+SEC across all three replicates, representing an approximate 21-fold increase (Figure 3M, Supplementary Table 2) and reflecting substantial EV enrichment following SEC. Proteomics was performed on the TFF-only and TFF+SEC EV samples to compare the types and distributions of protein identifications. Principal component analysis (PCA) plot of the protein levels showed clear separation between TFF-only and TFF+SEC preparations, indicating marked changes in the sample protein compositions after subsequent purification of the TFF-EVs by SEC (Figure 3N). The number of proteins that could be identified by mass spectrometry was substantially higher in the TFF+SEC replicates (607, 612 and 617 proteins) than in TFF-only (361, 339 and 364 proteins, 8.6 to 14.8% missingness, Figure 3P), consistent with improved EV sample purity following SEC. This was accompanied by improved reproducibility, with the CV% of protein intensities across the three technical replicates, decreasing from 36.8% for TFF-only versus to 12.4% for TFF+SEC (Figure 3O) and enrichment of integral EV proteins such as CD9, PDCD6IP and SDCBP (Figure 7A).

**Figure 7:**
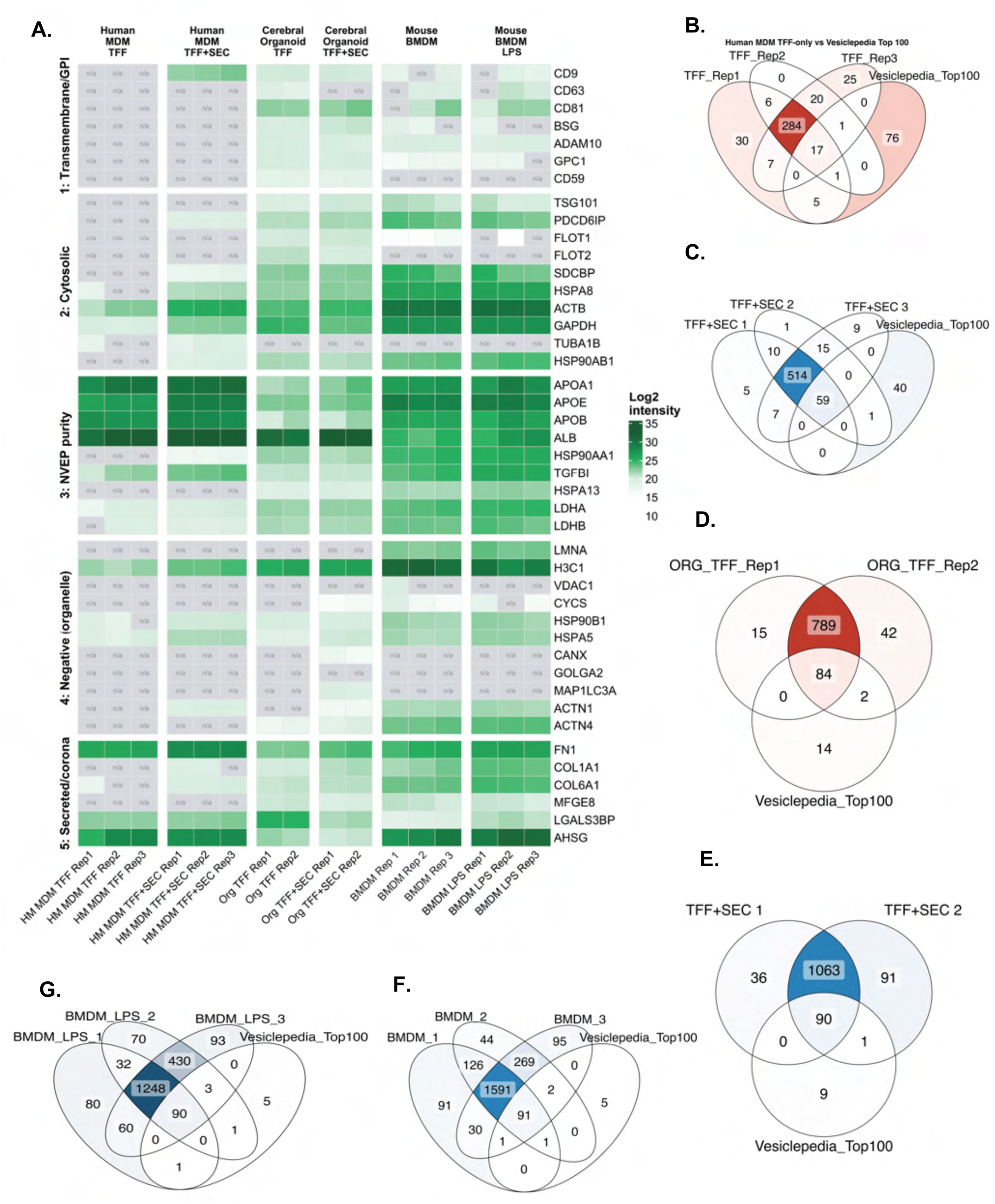
Protein marker validation against MISEV2023 guidelines and Vesiclepedia benchmarking across all three EV sources. (A) Heatmap of protein markers spanning the five MISEV2023 Table 3 categories (Welsh et al. 2024, *J Extracell Vesicles*) (1) across all six sample groups (human MDM TFF-only and TFF+SEC, human cerebral organoid TFF-only and TFF+SEC, mouse BMDM and BMDM_LPS), individual replicates shown as separate columns: • Category 1: Transmembrane/GPI-anchored proteins: CD9, CD63, CD81 (tetraspanins), BSG, ADAM10, GPC1, CD59 • Category 2: Cytosolic proteins: TSG101, PDCD6IP (ESCRT-associated), FLOT1, FLOT2, SDCBP (flotillin/syntenin), HSPA8, ACTB, GAPDH, TUBA1B, HSP90AB1 • Category 3: Non-vesicular extracellular particles (NVEPs): APOA1, APOE, APOB, ALB, HSP90AA1, TGFBI, HSPA13, LDHA, LDHB • Category 4: Organelle markers: LMNA, H3C1 (nucleus), VDAC1, CYCS (mitochondria), HSP90B1, HSPA5, CANX, GOLGA2 (ER/Golgi), MAP1LC3A (autophagosome), ACTN1, ACTN4 • Category 5: Secreted proteins/protein corona co-isolated with EVs: FN1, COL1A1, COL6A1 (ECM), MFGE8, LGALS3BP, AHSG For mouse BMDM samples, human marker proteins were mapped to their confirmed mouse orthologs. Cell colour indicates log2 protein intensity (DIA-NN); grey cells marked “n/a” indicate the protein was not detected in that replicate. (B-G) Overlap between proteins detected per biological replicate and the Vesiclepedia top-100 most frequently reported EV marker list (17), shown as Venn diagrams for (B) Human MDM TFF-only, (C) Human MDM TFF+SEC, (D) Cerebral Organoid TFF-only, (E) Cerebral Organoid TFF+SEC, (F) Mouse BMDM, and (G) Mouse BMDM_LPS preparations.

### Serum-free human cerebral organoid conditioned media shows minimal protein co-concentration during TFF with SEC purification

An identical protocol was applied to the serum-free CM from the human cerebral organoid. Due to limited sample availability, two technical replicates were processed (n=2). Permeate flow rate during sample processing was higher and required less clamp adjustment than for human MDM (15 to 16

drops/30 s versus 10 to 11 drops/30 s, Table 2), indicating lower foulant load. NTA showed TFF-only particle counts for the two replicates (1.39×10^10^ and 5.65×10^10^) were similar to the TFF+SEC counts (1.37×10^10^ and 1.22×10^10^; Figure 4A-B, E-F). However, as with human MDM, particles in the TFF+SEC EV preparations had a narrower, more consistent size distribution than TFF-only, with mode ranging from 87.5 to 117.5 nm (mean 102.5 ± 21.2 nm) and zetaview cumulative particle population at D1-D90 spanning roughly 70 to 235 nm for TFF-only preparations, compared to a narrower mode ranging from 117.5 to 137.5 nm (mean 127.5 ± 14.1 nm) and D10-D90 spanning roughly 80 to 235 nm for TFF+SEC (Figure 4A-B, E-F). Notably, for TFF+SEC preparations, the CMG membrane-positive fraction was markedly lower in cerebral organoid CM preparations (5.9% ± 4.9%) than in human MDM (35.4% ± 11.9%), exhibiting a marked difference in its composition.

Moreover, although EVs from cerebral organoid CM showed recovery of integral EV proteins (CD9, CD81 TSG101, PDCD6IP) for both TFF-only and TFF+SEC preparations, the TFF-only preparation still had a protein concentration of 378-984 µg/ml which substantially reduced for the TFF+SEC to 52.7-70µg/ml (Supplementary Table 1). For the TFF-only preparation from cerebral organoids CM, TEM grids contained identifiable vesicular structures from the cerebral organoids (Figure 4C, G) and did not contain a dense protein background that was observed for the human MDM TFF-only preparations. TFF+SEC preparations from cerebral organoid CM imaged by TEM revealed even cleaner examples of EVs adsorbed to the grids, with minimal background (Figure 4D, H). In cerebral organoid CM, particle-to-protein ratio of the replicates increased from 2.13×10^8^ and 2.01×10⁷ particles/µg for TFF-only preparations to 4.34×10^8^ and 6.51×10^8^ particles/µg (i.e. ∼2-to 32-fold) for TFF+SEC preparations (Figure 4I, Supplementary Table 2). Notably, organoid TFF-only preparations already had a particle-to-protein ratio ∼10.7-fold higher on average than TFF-only preparations from human MDM CM (1.09×10⁷ particles/µg, Supplementary Table 2). More proteins were identified per replicate following SEC (1200 and 1258) than in TFF-only preparations (895 and 924, Figure 4K), consistent with the pattern seen in human MDM and likely reflecting depletion of dominant abundant proteins, allowing lower-abundance proteins to be detected with greater resolution. In addition, PCA of the protein levels showed clear separation between the two conditions in regard to detected protein composition and level (Figure 4J). However, while the TFF system alone performs well for isolating comparatively clean EVs from serum-free organoid conditioned media, subsequent SEC further altered the protein profile as indicated by PCA. It is important to note, while the media for organoid culture had no serum supplemented, there may be proprietary additions of some level of serum factors as we detected albumin by proteomics in both TFF only and TFF+SEC preparations (Figure 7A).

### Differentially abundant proteins following SEC are enriched for distinct functional categories in each condition

Next, we wanted to further assess the difference in protein components between the samples before and after SEC cleanup post TFF. Differential expression analysis identified 71 of 342 testable proteins significantly higher in abundance TFF-only preparations and 60 significantly higher in TFF+SEC preparations for human MDM (adj p-val < 0.05; Figure 5A). For cerebral organoid CM preparations, we identified 116 of 872 testable proteins significantly higher in TFF-only and 93 significantly higher in TFF+SEC preparations (adj p-val < 0.05; Figure 5D).

In human MDM-derived CM, proteins significantly higher in TFF-only preparations were enriched for immunoglobulin-mediated immune response (24 proteins), complement activation (5 proteins), extracellular region (67 proteins), antigen binding (27 proteins), and Toll-like receptor 4 binding (2 proteins) (Figure 5B). Proteins significantly higher in TFF+SEC were enriched for blood coagulation (17 proteins), protein maturation (18 proteins), extracellular region (57 proteins), cell adhesion molecule binding (18 proteins), and extracellular matrix structural constituent (12 proteins) (Figure 5C).

In cerebral organoid CM, proteins significantly higher in TFF-only preparations were enriched for humoral immune response (16 proteins), fibrinolysis (6 proteins), extracellular exosome (76 proteins), endomembrane system (57 proteins), endopeptidase inhibitor activity (11 proteins), and structural molecule activity (22 proteins) (Figure 5E). Proteins significantly higher in TFF+SEC preparations were enriched for cell migration (26 proteins), nervous system development (31 proteins), extracellular vesicle (54 proteins), synapse (33 proteins), cell adhesion molecule binding (16 proteins), and enzyme binding (25 proteins) (Figure 5F). Full protein lists for these GO-enriched groups are provided in Supplementary Table 9.

To further characterise proteins specifically associated with TFF-only preparations, we identified proteins detected in every TFF-only technical replicate, that were absent from every replicate of TFF+SEC, for human MDM and cerebral organoid (Supplementary Figure 3). This presence/absence classification identified 21 TFF-only-unique proteins in human MDM consisting of mostly histones and immunoglobulins and 63 in cerebral organoid consisting of various signalling molecules (see Supplementary Table 8). STRING 12.0 protein-protein interaction networks of these TFF-only-unique proteins are shown in Supplementary Figure 3, with full gene identity and mapping provided in Supplementary Table 8.

### The TriPort-TFF protocol is reproducible across biological replicates and distinguishes two biological conditions in mouse BMDM

Additionally, to test whether the system could support EV-based comparisons between biologically distinct experimental groups, we applied the protocol to CM from mouse bone marrow-derived macrophages (BMDM), unstimulated or stimulated with LPS (BMDM_LPS, 1µg/ml for 24-h), using three independent biological replicates (individual animals) per condition (n=3, Figure 6). Peak particle concentration varied approximately three-fold between replicates within each group (Figure 6A, C), consistent with expected biological variability between animals. Representative TEM confirmed EV morphology in both BMDM and BMDM_LPS preparations (Figure 6B, D).

At the protein level, PCA of protein intensities showed clear separation between BMDM and BMDM_LPS (PC1 = 46.9%, PC2 = 23.4% of the total variance; Figure 6E), demonstrating distinct EV proteomic differences between the two conditions. Inter-replicate reproducibility was high for both biological conditions, reflected by low CV% (Figure 6F), with a median of 23% for BMDM and 21.8% for BMDM_LPS. Protein identifications averaged 2052 ± 101 proteins for BMDM (range 1939-2131) and 1776 ± 225 proteins for BMDM_LPS (range 1516-1931; Figure 6G).

### EVs isolated via the lab-assembled TFF system show source- and purification-dependent differences in canonical marker detection

To characterize the composition of isolated particles, we assessed detection of protein markers spanning the five MISEV2023 Table 3 categories: Category 1, transmembrane/GPI-anchored proteins; Category 2, cytosolic proteins recovered in EVs; Category 3, non-EV co-isolated structures used as purity controls; Category 4, intracellular compartment markers used as negative controls; and Category 5, secreted proteins recovered with EVs (1) (Figure 7A). Canonical transmembrane and cytosolic EV markers were largely undetected in human MDM TFF-only EV preparations but became detectable by LC-MS/MS following SEC, including CD9, PDCD6IP (ALIX), SDCBP, HSPA8, ACTB, GAPDH, and HSP90AB1. Cerebral organoid and mouse BMDM preparations showed more consistent detection of Category 1 and 2 markers across both TFF-only and TFF+SEC conditions. CD59, the sole Category 1c marker included, was detected only in organoid preparations and was absent from both human MDM and mouse BMDM (Figure 7A). Humans carry a single CD59 gene, whereas mice carry two functionally distinct paralogues (Cd59a, Cd59b) (18), neither of which was detected in BMDM condition.

Next, to assess protein detection consistency across replicates, proteins identified in each replicate were compared against the Vesiclepedia top-100 EV marker list, a manually curated, community-annotated database of proteins reported in extracellular vesicles across published studies (17) (Figure 7B-G). Overlap with this reference set was modest across all three sources (5.4-10.3% of core detected proteins) and increased following SEC for both human MDM EVs (17/100 TFF-only; 59/100 TFF+SEC) and cerebral organoid EVs (84/100 TFF-only; 90/100 TFF+SEC). A comparable proportion of core proteins overlapped with the reference set in mouse BMDM EVs under both conditions (91/100, BMDM; 90/100, BMDM_LPS). Full protein lists for each Venn category are provided in Supplementary Table 4.

## Discussion

EV isolation methods can be limited by the nature of their relatively low concentrations in high volumes of samples including biological and CM fluids. The system validated here, TriPort-TFF, named for its three-port cap assembly, is a peristaltic pump-driven, hollow-fiber TFF setup built entirely from individually sourced components (Table 1), centered on a custom cap assembly with needle and stopcock ports, and a 500 kDa Repligen MicroKROS membrane. Every operating parameter (flow rate, pressure inference via droplet counting, cleaning and retirement criteria) is specified in the Methods allowing the system to be assembled at relatively low cost (see Supplementary Table 6). The workflow detailed here involved TFF alone and combined with SEC preparations, which reproducibly isolated EVs from serum-containing CM of human MDM and mouse BMDM and from serum-free CM of human cerebral organoid cultures. CM from LPS-stimulated and unstimulated BMDM were compared, revealing distinct proteomic clusters in EVs isolated from each condition.

Since this system has no integrated pressure gauge, TMP was inferred from permeate flux, calculated from droplet count and droplet volume (LMH). Rather than using a conventional water flux recovery (WFR) metric (8), a standard metric for baseline membrane performance, the membrane condition was tracked using PBS Flux Recovery (PFR), since PBS buffer was used for all the samples. Although this is a coarser method that cannot directly detect transient fine pressure changes, it is a trade-off for an accessible low-cost system. Labs that require finer control could modify the system to add an inline pressure monitor. Further, the cap assembly design is not necessarily restricted to a 50 mL falcon tube, the same construction principle could be applied to a larger container for higher input volumes, though this was not tested in the present study.

Beyond confirming system integrity, permeate flow rate during sample processing also carried information about the sample itself. Once a sample was introduced, flow rate dropped from the pristine-membrane priming range and stabilized at source-dependent levels: 10-11 drops/30 seconds for human MDM, 11-12 for mouse BMDM, and 15-16, closer to the priming baseline, for cerebral organoid (Table 2). The lower stabilized flow rates for the two serum-containing CM sources relative to serum-free organoid CM likely reflected a higher foulant load building up on the membrane during processing of fluids containing serum proteins.

The capacity of the TFF protocol to isolate reproducible EV populations varied meaningfully depending on the source of the CM and the incorporation of SEC into the workflow. For CM from human MDM, inter-replicate CV dropped from a median of 36.8% after TFF-only processing, to 12.4% after TFF+SEC processing of the CM. For cerebral organoid CM, the CV was not considered a reliable indicator at n=2 replicates, but on a descriptive basis, protein detection counts differed by only 3-5% between the two replicates in each isolation condition (895-924 proteins in TFF-only and 1200-1258 in TFF+SEC).

For CM from unstimulated or LPS-treated mouse BMDM cultures, biological replicates from independent animals showed a CV of 23% (BMDM) and 21.8% (BMDM_LPS), with 1682 and 1338 proteins shared across all three replicates of each condition respectively (Supplementary Table 4). This dataset also shows that the system preserves biologically meaningful separation between conditions, not just technical consistency. A TFF-only versus TFF+SEC preparation comparison for BMDM was also assessed separately in pilot run (Supplementary Figure 2 and 4) to confirm enrichment of EVs after SEC processing. Taken together, these results indicate that reproducibility depends on sample type, replicate number, and, in particular for serum-containing sources of CM, inclusion of a purification step like SEC in the workflow.

For serum-containing CM preparations (human MDM, mouse BMDM), TFF alone concentrated substantial non-EV/ serum protein alongside EVs, as protein concentration in TFF-only preparations from human MDM CM ranged from 4500-5788 µg/ml, compared to 75.2-80.4 µg/ml following SEC (Supplementary Table 1). TEM showed a dense, heterogenous protein background in TFF-only preparations that was largely resolved after SEC. SEC also improved purity, with the particle-to-protein ratio increasing ∼21-fold from TFF-only to TFF+SEC preparations.

Interestingly, in TFF+SEC preparations from CM of human MDM, enriched proteins represented blood coagulation pathway related proteins, including fibrinogen chains, complement factor C4BPB, and Apolipoprotein E (Supplementary Table 9). These proteins have previously been reported as present in the formation of the EV corona (19), a physiologically associated layer of plasma-derived protein on the surface of the EV, which in this culture system could potentially be from EV-depleted autologous human serum supplement. In this study, we cannot distinguish corona-associated protein from residual free plasma protein using the present data alone without an orthogonal approach for corona stripping and analysis. Albumin, on the other hand, was detected across all sample groups regardless of the source or purification steps. SEC processing post TFF of CM from Human MDM, however, largely removed nuclear proteins (H2BC4, H2BC5, H2BC7, H2BC9, H2BC13, H2BC14, H2BC15, H2BC18). We also note that “extracellular region” is a broad, non-specific GO:CC functional pathway term that was independently top-ranked on both TFF-only and TFF+SEC preparations of human MDM comparison, limiting how cleanly this enrichment analysis alone can separate contaminant from EV-associated signal (protein lists are provided in Supplementary Table 9).

For CM from mouse BMDM, TFF-only preparation protein content was similarly high relative to the downstream TFF+SEC-purified preparation (203 µg of total protein pre-SEC, pilot (n=1), see Supplementary Figure 2C and 4), consistent with the same serum-protein co-concentration pattern observed in preparations from CM of human MDM. Notably, despite both the mouse and human CM containing serum, TFF retentate from CM of human MDM containing EV-depleted autologous human serum was visibly yellow (Supplementary Figure 1B), while TFF retentate from CM of BMDM (EV-depleted FBS) was not (Supplementary Figure 2B). We speculate that this reflects serum pigmentation, possibly bilirubin bound to albumin, an established yellow-orange chromophore in human serum (20), though this was not directly investigated.

For serum-free human organoid CM, TFF alone yielded a ∼10.7-fold higher particle-to-protein ratio and cleaner morphology of EVs imaged by TEM than the serum-containing sources CM derived preparations. As with CM of human MDM, protein concentrations were higher in TFF-only preparations from organoid CM (378-984 µg/ml) compared to TFF+SEC preparations (52.7-70 µg/ml, Supplementary Table 1), although the overall serum-protein concentrations were lower in organoid CM than in preparations from CM of human MDM (4500-5788 µg/ml). Despite this relatively cleaner starting point, we cannot conclude that SEC in organoid preparations serves only to remove residual contaminants. Proteins specifically detected in TFF-only preparations from organoid CM were not dominated by obvious contaminants in the way the CM of human MDM TFF-only-unique set was mostly histones and immunoglobulins (Supplementary Figure 3). The cerebral organoid CM was pooled from day 90, 93, 96 and 100, at which stages organoids would have superficial cortical neurons and be starting to undergo gliogenesis (15). Consistent with this developmental window, the 63 proteins detected exclusively in TFF-only preparations from CM of organoid culture, include HPCA (hippocalcin), a neuronal calcium-sensor protein, alongside adhesion and signaling proteins such as CD44 and SDC1 (Supplementary Figure 3). This set also included NFIA (Nuclear Factor I A), a transcription factor previously identified as a driver of the neurogenesis-to-gliogenesis transition in stem-cell and organoid models (21,22). Cerebral organoid derived-EVs have separately been shown to carry cell-type-specific and developmentally dynamic protein cargo (23), so whether the specific proteins identified here represent functional cargo lost upon SEC, rather than non-specific co-isolate, cannot be resolved from proteomic network data alone and would require further cell-type-of-origin or functional validation.

This ambiguity is also reflected at the pathway level where proteins significantly higher in organoid TFF-only compared to TFF+SEC preparations were enriched for GO:CC term “extracellular exosome,” suggesting loss of some EV-associated markers after SEC. Conversely, proteins significantly higher in organoid TFF+SEC were enriched for “nervous system development” and synapse terms specifically alongside “extracellular vesicles,” suggesting that TFF+SEC may reveal tissue specific, central nervous system (CNS) relevant EV biology that is otherwise masked in TFF-only preparation from organoid CM. Additionally, several Category 4 negative markers, including CYCS (mitochondrial), CANX (ER), and MAP1LC3A (autophagosome), were undetected in organoid TFF-only preparation but became detectable following SEC. A similar low-abundance unmasking pattern was observed for canonical EV-positive markers in CM from human MDM cultures, suggesting SEC increases proteomic sensitivity broadly across sample types.

Consistent with the particle-to-protein and proteomic depth improvements described above, SEC purification was accompanied by a marked increase in Vesiclepedia marker recovery in Human MDM (17 to 59) and a smaller increase in cerebral organoid preparations (84 to 90). BMDM preparations, assessed here as TFF+SEC only, showed comparably high marker recovery (90–91), consistent with the pattern observed in the other two sources. Given the small, fixed size of this reference list, and that its composition reflects frequency of reporting across the broader EV literature rather than these specific sources, we treat this comparison as one general cross-check, alongside the MISEV2023 category framework.

Several limitations should be considered in interpreting these results. The CM from human MDM cultures were derived from a single donor; the n=3 technical replicates demonstrate protocol-level reproducibility from a shared input but cannot account for donor-to-donor variability, which remains untested here for human MDM. The organoid dataset (n=2) was constrained by sample availability and should be read as a proof-of-concept demonstration of EV isolation from CM of these cultures. This study has tested the TFF system on CM only; performance on other complex fluids such as urine or milk was not assessed.

In conclusion, we have demonstrated that the TriPort-TFF workflow detailed here reproducibly isolates EVs from CM of different cell culture systems, and it is sensitive enough to detect a biological effect in CM of mouse BMDM culture treated with LPS. Whether SEC is a necessary addition to TFF for isolation and analysis of EVs depends on the complexity of the fluid, for example serum content of the input medium as well as the specific biological question being asked. The workflow and results presented here provides researchers with a practical basis for adapting this protocol to their own sample type rather than defaulting to always or never including a polishing step. We offer this system as an accessible, transparently validated reference protocol rather than a competitive alternative to industrial-grade systems, in the hope of lowering the practical barrier for TFF adoption to support broader, more standardized use of this isolation method across the EV research community.

## Abbreviations

BMDM: Bone marrow-derived macrophages
CM: Conditioned Media
CV: Coefficient of Variation
EV: Extracellular Vesicles
FBS: Fetal bovine serum
GO:CC: Gene Ontology: Cellular Component
LMH: Liters per square meter per hour (flux unit)
LPS: Lipopolysaccharide
MDM: Monocyte-derived macrophage
MISEV: Minimal Information for Studies of Extracellular Vesicles
NTA: Nanoparticle tracking analysis
PFR: PBS Flux Recovery
SEC: Size-exclusion chromatography
TEM: Transmission Electron Microscopy
TFF: Tangential Flow Filtration
TMP: Transmembrane pressure
WFR: Water Flux Recovery

## Acknowledgements

We thank Sydney Mass Spectrometry, the Sydney Microscopy & Microanalysis Facility, and Sydney Analytical (core research facilities at the University of Sydney) for technical support and access to instrumentation. D.M.’s research visit to the laboratory of Dr Setty Magaña at The Abigail Wexner Research Institute at Nationwide Children’s Hospital, supported by the Ian Ballard Travel Award (MS Australia), provided foundational training in the EV isolation and characterization techniques underlying this study. Diagrams were created with BioRender.com.

## Data Availability Statement

All data supporting the findings of this study are included in the manuscript and its Supplementary Information. Any additional data are available from the corresponding author upon request.

