## Supplementary File 2 (Figures 1-4 and Table 7) for "TriPort tangential flow filtration: a lab-assembled system for EV isolation across diverse cell-culture sources"

File 1

**Supplementary Figure 1.** Pilot study of Human MDM TFF-only retentate across individual SEC elution fractions (F1-F4), showing reference marker recovery, protein yield, marker intensity, and fraction overlap.

**Supplementary Figure 2.** Pilot study of Mouse BMDM TFF-only retentate across individual SEC elution fractions (F1-F4), showing reference marker recovery, protein yield, marker intensity, and fraction overlap.

**Supplementary Figure 3.** Shared and TFF-only-unique proteins between technical replicates for Human MDM and Cerebral Organoid, with STRING protein-protein interaction networks of the TFF-only-unique proteins.

**Supplementary Table 7.** Reference protein sets used for EV marker recovery quantification in Supplementary Figs. 1A and 2A.

**Supplementary Figure 4.** Transmission Electron Microscope micrographs of EVs isolated from CM of Human MDM, Cerebral organoid, BMDM and liposomes at 500nm scale.

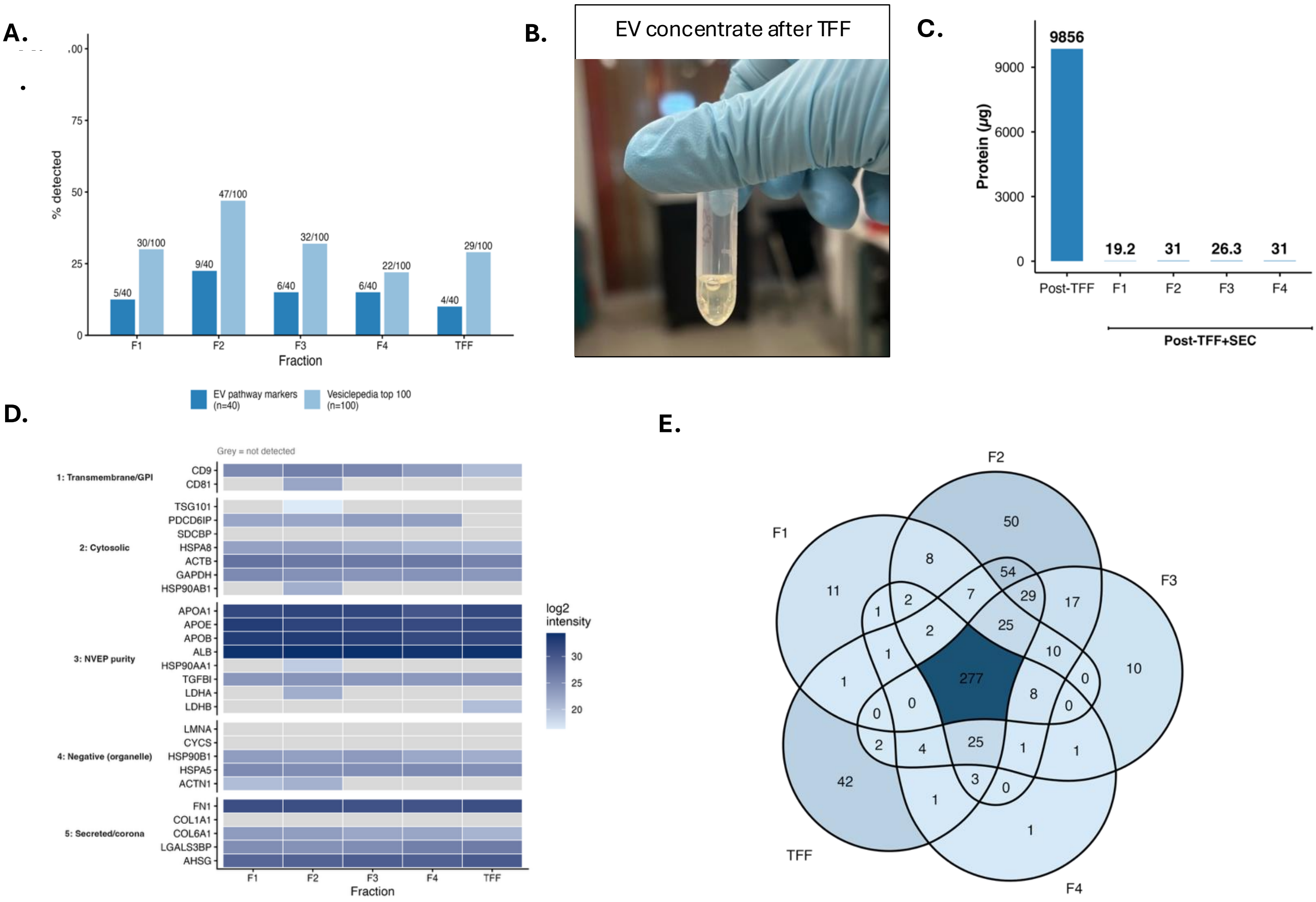


**Supplementary Figure 1. Pilot study of Human MDM TFF-only retentate across individual SEC elution fractions (F1-F4), showing reference marker recovery, protein yield, marker intensity, and fraction overlap.** Pilot run was performed prior to the replicate experiments reported in the main text. This was used to characterize EV marker recovery across individual SEC elution fractions and confirm the necessity of SEC as a downstream purification step. (A) Percentage of two reference marker sets detected in the TFF-only retentate (pre-SEC comparator) and in each individual SEC elution fraction (F1-F4): Vesiclepedia top 100 (dark blue) and EV pathway markers, n=40 (light blue), spanning four defined EV biogenesis mechanisms plus a caveated abundant-cytosolic-protein category (Supplementary Table 7 below). Labels indicate the absolute count and percentage of each reference set detected per fraction. (B) Representative photograph of the concentrated EV retentate immediately following TFF processing, prior to SEC; the yellow coloration reflects residual serum pigment carried over from the serum-containing conditioned media. (C) Total protein yield (µg) in the TFF-only retentate and in each SEC elution fraction, quantified by Qubit fluorometric protein assay. (D) Heatmap of log2 DIA intensity shows protein markers spanning the five MISEV2023 Table 3 categories, across the TFF-only retentate and each SEC fraction: transmembrane/GPI-anchored proteins (Category 1: CD9, CD63, CD81, BSG, ADAM10, GPC1, CD59), cytosolic proteins recovered in EVs (Category 2: TSG101, PDCD6IP, FLOT1, FLOT2, SDCBP, HSPA8, ACTB, GAPDH, TUBA1B, HSP90AB1), non-EV co-isolated structures used as purity controls (Category 3: APOA1, APOE, APOB, ALB, HSP90AA1, TGFBI, HSPA13, LDHA, LDHB), intracellular compartment negative markers (Category 4: LMNA, H3C1, VDAC1, CYCS, HSP90B1, HSPA5, CANX, GOLGA2, MAP1LC3A, ACTN1, ACTN4), and secreted proteins/protein corona (Category 5: FN1, COL1A1, COL6A1, MFGE8, LGALS3BP, AHSG). Grey tiles indicate protein not detected in that fraction. (E) Venn diagram showing the number of proteins detected exclusively in the TFF-only retentate versus each SEC fraction, and shared across combinations (TFF, F1, F2, F3, F4).

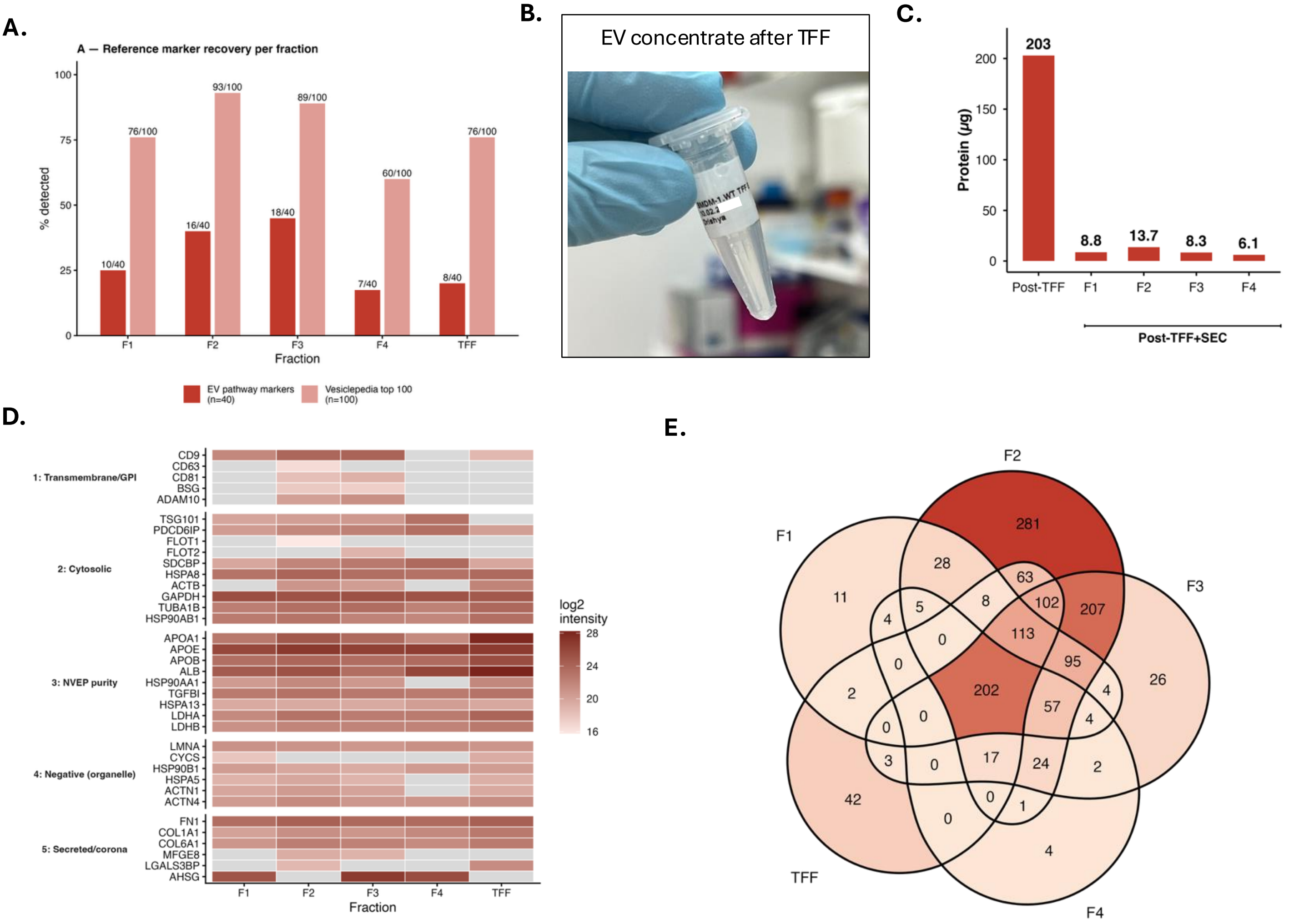


**Supplementary Figure 2. Pilot study of Mouse BMDM TFF-only retentate across individual SEC elution fractions (F1-F4), showing reference marker recovery, protein yield, marker intensity, and fraction overlap.** Pilot run was performed prior to the replicate experiments reported in the main text, used to characterize EV marker recovery across individual SEC elution fractions and confirm the necessity of SEC as a downstream purification step. (**A**) Percentage of two reference marker sets detected in the TFF-only retentate (pre-SEC comparator) and in each individual SEC elution fraction (F1-F4): Vesiclepedia top 100 (dark red) and EV pathway markers, n=40 (orange), spanning four defined EV biogenesis mechanisms plus a caveated abundant-cytosolic-protein category (Supplementary Table 7 below). Labels indicate the absolute count and percentage of each reference set detected per fraction. (**B**) Representative photograph of the concentrated EV retentate immediately following TFF processing, prior to SEC. (**C**) Total protein yield (µg) in the TFF-only retentate and in each SEC elution fraction, quantified by Qubit fluorometric protein assay. (**D**) Heatmap of log2 DIA intensity shows protein markers spanning the five MISEV2023 Table 3 categories, across the TFF-only retentate and each SEC fraction: transmembrane/GPI-anchored proteins (Category 1: CD9, CD63, CD81, BSG, ADAM10, GPC1, CD59), cytosolic proteins recovered in EVs (Category 2: TSG101, PDCD6IP, FLOT1, FLOT2, SDCBP, HSPA8, ACTB, GAPDH, TUBA1B, HSP90AB1), non-EV co-isolated structures used as purity controls (Category 3: APOA1, APOE, APOB, ALB, HSP90AA1, TGFBI, HSPA13, LDHA, LDHB), intracellular compartment negative markers (Category 4: LMNA, H3C1, VDAC1, CYCS, HSP90B1, HSPA5, CANX, GOLGA2, MAP1LC3A, ACTN1, ACTN4), and secreted proteins/protein corona (Category 5: FN1, COL1A1, COL6A1, MFGE8, LGALS3BP, AHSG). Grey tiles indicate protein not detected in that fraction. (**E**) Venn diagram showing the number of proteins detected exclusively in the TFF-only retentate versus each SEC fraction, and shared across combinations (TFF, F1, F2, F3, F4).

| EV Pathway markers (n=40) | Vesiclepedia top 100 |
|---|---|
| TSG101, PDCD6IP, VPS4A, VPS4B, HRS, STAM, EAP20, SNF8, VPS25, CHMP4A, CHMP4B, CHMP4C, CHMP2A, CHMP2B, CHMP3, CHMP1A, CHMP1B, CHMP6, IST1, SDCBP, SDC1, SDC2, SDC3, SDC4, SMPD3, FLOT1, FLOT2, RAB31, CD63, CD9, CD81, CD82, CD151, TSPAN8, TSPAN4, HSPA8, HSP90AA1, HSP90AB1, ACTB, GAPDH | CD63, CD9, PDCD6IP, TSG101, CD81, GAPDH, FLOT1, ACTB, ANXA2, SDCBP, HSP90AA1, HSPA8, ANXA5, ENO1, PKM, HSP90AB1, YWHAZ, PGK1, YWHAE, FLOT2, ANXA1, VCP, PPIA, EEF1A1, ALB, ALDOA, ITGB1, MYH9, CLTC, TPI1, CFL1, EEF2, MSN, GNB1, PRDX1, ANXA6, SLC3A2, EZR, LDHA, LDHB, BSG, CDC42, PFN1, ATP1A1, ACTN4, HSPA1A, FLNA, YWHAB, GNAI2, YWHAQ, FASN, CLIC1, PRDX2, GSN, CCT2, RAB5C, HIST1H4A, RAP1B, GNB2, LGALS3BP, YWHAG, RAB10, HLA-A, ACTN1, ANXA7, FN1, TFRC, RAN, GDI2, CCT3, AHCY, HSPA5, CCT4, ACLY, C3, UBA1, ANXA11, TUBB4B, KPNB1, CAP1, RAC1, MFGE8, TCP1, RHOA, TLN1, CCT6A, GNAS, CCT5, RALA, EHD1, CCT8, PGAM1, IQGAP1, VCL, GPI, EIF4A1, RAB7A, EEF1G, ADAM10, A2M |

**Supplementary Table 7. Reference protein sets used for EV marker recovery quantification in Supplementary Figs. 1A and 2A.** Two fixed reference protein sets were used to quantify EV marker recovery per fraction. EV Pathway markers (n=40) is a curated, mechanism-based set spanning four defined EV biogenesis pathways: ESCRT machinery (TSG101, PDCD6IP, VPS4A/B, HRS/STAM, CHMP2A/2B/3/4A/4B/4C/6, IST1), the Syndecan-Syntenin-ALIX pathway (SDCBP, SDC1-4), the ceramide/RAB31-flotillin pathway (FLOT1, FLOT2, RAB31, SMPD3), and tetraspanins (CD63, CD9, CD81, CD82, CD151, TSPAN4, TSPAN8), plus a fifth category of abundant cytosolic proteins associated with EVs (HSPA8, HSP90AA1, HSP90AB1, ACTB, GAPDH). Vesiclepedia top 100 is the 100 most frequently detected EV proteins in the Vesiclepedia database, used as a broad, literature-derived EV proteome reference independent of any specific biogenesis mechanism.

References for EV Pathway markers: MISEV2023 (Welsh et al., Journal of Extracellular Vesicles 2024) (1). Vesiclepedia top 100: Chitti et al., Nucleic Acids Research 2024 (17).

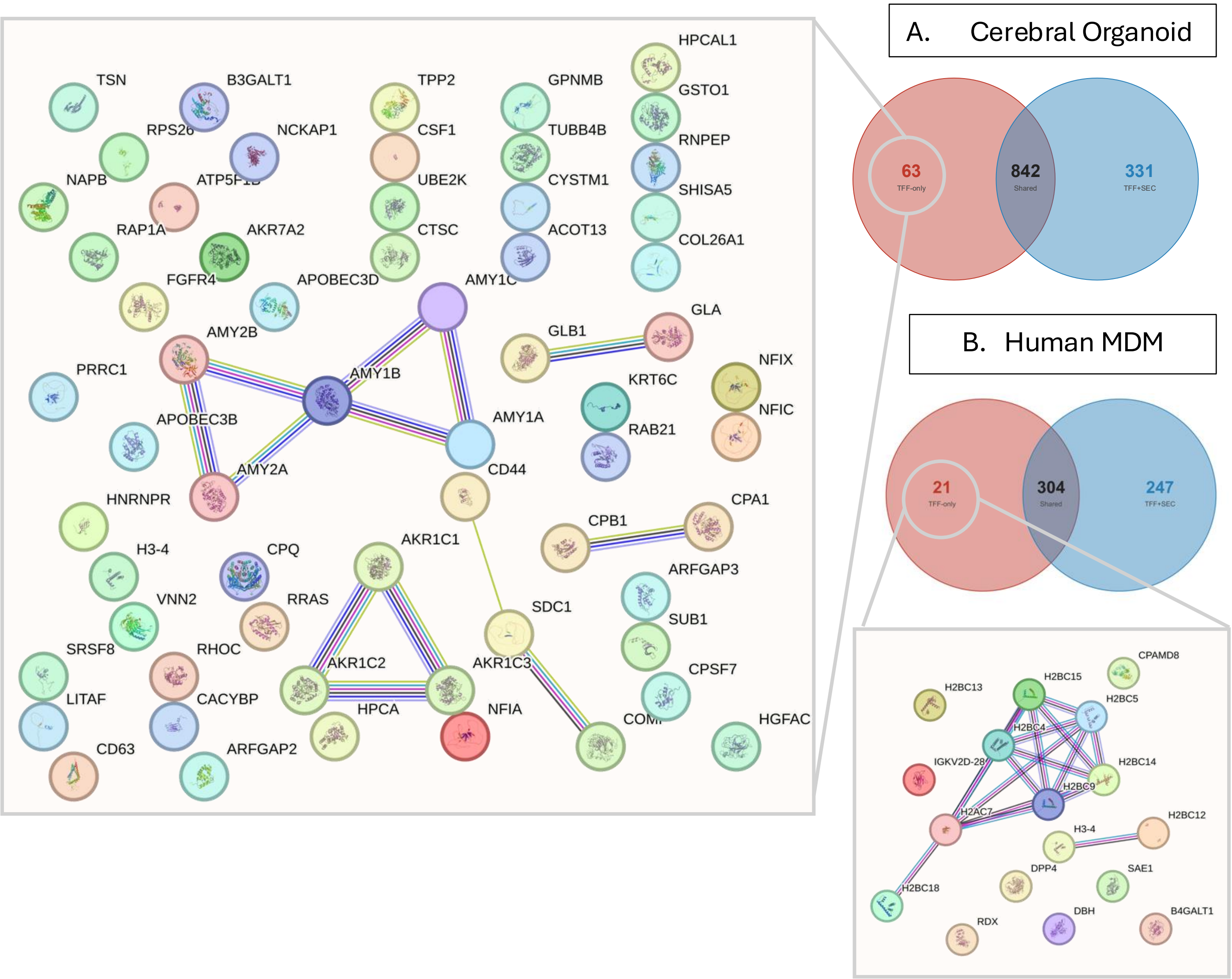


**Supplementary Figure 3. Shared and TFF-only-unique proteins between technical replicates for Human MDM and Cerebral Organoid, with STRING protein-protein interaction networks of the TFF-only-unique proteins.**

(A) Venn diagram showing proteins detected in all replicates of Cerebral Organoid TFF-only, all replicates of TFF+SEC, or both: 63 proteins detected in all TFF-only replicates and absent from all TFF+SEC replicates, 331 proteins detected in all TFF+SEC replicates and absent from all TFF-only replicates, and 842 proteins detected in all replicates of both conditions. STRING protein-protein interaction network of the 63 proteins detected exclusively in Cerebral Organoid TFF-only preparations (inset, left). (B) Venn diagram showing the same comparison for Human MDM: 21 proteins detected in all TFF-only replicates and absent from all TFF+SEC replicates, 247 proteins detected in all TFF+SEC replicates and absent from all TFF-only replicates, and 304 proteins detected in all replicates of both conditions. STRING protein-protein interaction network of the 21 proteins detected exclusively in Human MDM TFF-only preparations (inset, bottom right). A protein was classified as condition-unique if detected in every replicate of one condition and no replicate of the other.

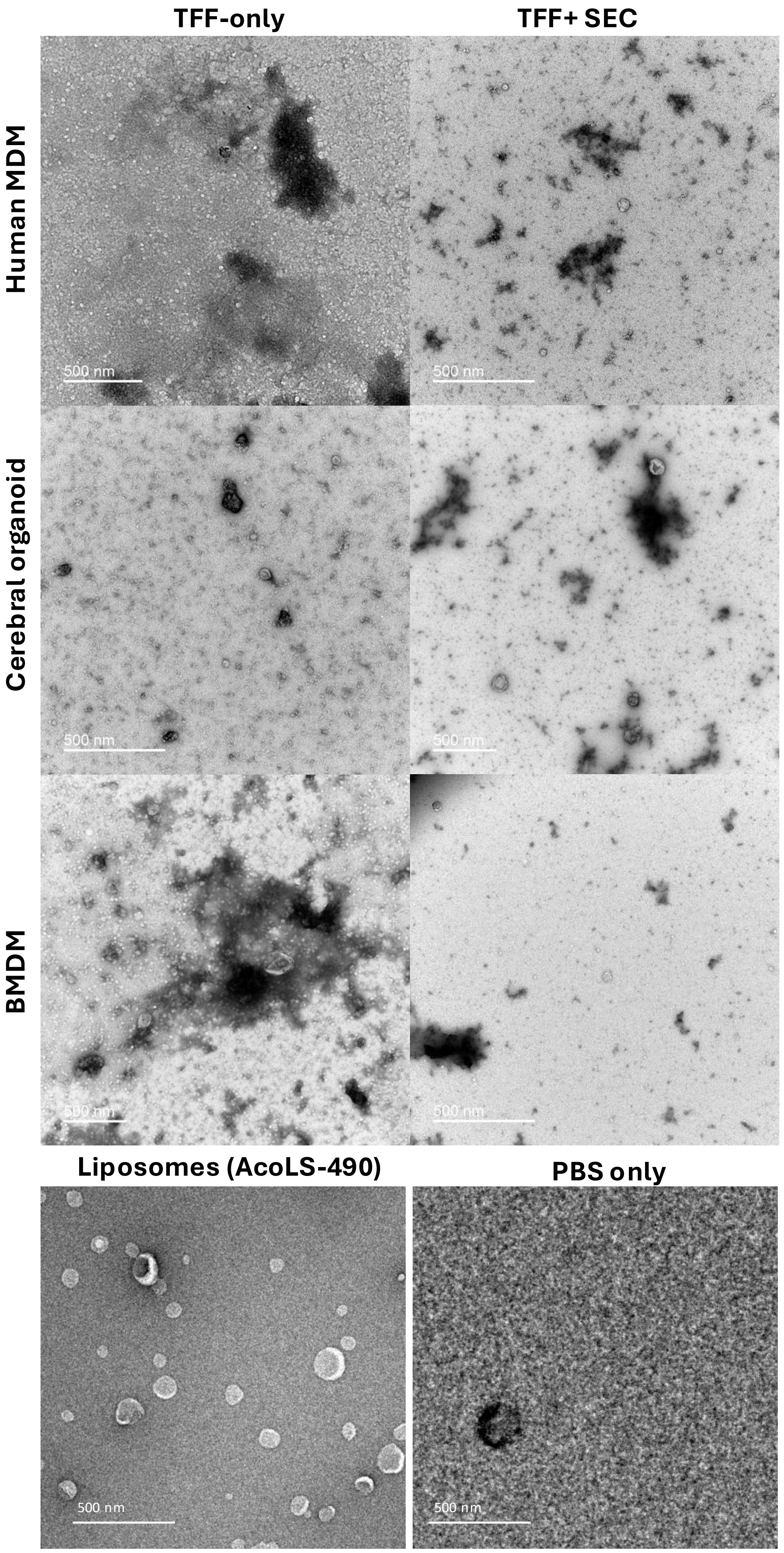
TFF-only
TFF+ SEC
Human MDM
Cerebral organoid
BMDM
500 nm
Liposomes (AcoLS-490)
PBS only

**Supplementary Figure 4. Transmission Electron Microscope micrographs of EVs isolated from CM of Human MDM, Cerebral organoid, BMDM and liposomes at 500nm scale.**
Transmission electron microscopy (TEM) micrographs are shown at a wider field of view (500 nm scale bar) to provide additional transparency for each condition. TFF-only (left column) and TFF+SEC (right column) preparations are compared across three conditioned-media (CM) sources: Human MDM (top row), cerebral organoid (second row), and BMDM (third row). A liposome preparation (5ul from 1:10 dilution with PBS of AcoLS-490 liposomes 200nm lot np. 49023001, Acoerela) is included as a morphology reference, illustrating canonical bilayer-bound vesicular structures for visual comparison against the more heterogeneous background observed in biological EV preparations; this is a qualitative reference field and not a quantified comparison. A PBS-only (buffer, no sample) field is included as a negative control.
